# Sensory neuronal mTORC1 signaling establishes neuroimmune memory that initiates allergic immunity

**DOI:** 10.1101/2025.11.10.687723

**Authors:** Xueping Zhu, Haibo Yang, Haoting Zhan, Isabela J. Kernin, Ngoc Le, Dean R. Buttaci, Cai Han, Parth R. Naik, Lukas M. Altenburger, Elena Wu, Peri R Matatia, Sarah Zaghouani, Neal P. Smith, Rebecca Londoner, Cameron H. Flayer, Zhengwang Sun, Alexandra-Chloe Villani, Rod A Rahimi, Caroline L Sokol

## Abstract

Environmental allergens are enriched in protease activity, a feature that directly activates cutaneous sensory neurons, triggering itch and Substance P release, which promotes Th2-skewing CD301b^+^ dendritic cell migration and initiates allergic immunity. However, allergens are typically encountered as repeated, subthreshold exposures, and how these cumulatively induce sensitization is unknown. We identify a sensory neuron-intrinsic mechanism of neuroimmune memory that enables this process. Initial activation by protease allergens induces sustained mTORC1 signaling and PGC-1α-associated mitochondrial remodeling, establishing a metabolically primed state that amplifies neuroimmune responses upon allergen re-exposure. This heightened state, independent of adaptive immunity, drives enhanced itch, increased CD301b^+^ dendritic cell migration, and augmented Th2 differentiation. Disrupting neuronal mTORC1 signaling or mitochondrial stability abrogates this amplification while leaving primary responses intact. Importantly, this mechanism generalizes across distinct protease allergens, revealing mTORC1-driven metabolic reprogramming in sensory neurons as a form of innate neuroimmune memory underlying allergen cross-sensitization and polysensitization.

## INTRODUCTION

The type 2, or allergic, immune response is characterized by CD4^+^ Th2 differentiation and IgE production, and is triggered by a diverse stimuli, including helminths, venoms, and environmental allergens (Iwasaki and Medzhitov, 2015; Pritchard et al., 2021; Rahimi and Sokol, 2022). A unifying feature of many such stimuli is protease activity (Aalberse, 2000; Soh et al., 2023; Traidl-Hoffmann et al., 2009). We and others have shown that protease allergens are directly detected by TRPV1^+^ peptidergic sensory neurons in the skin, which elicit acute itch and release the neuropeptide substance P (SP) to promote mast cell activation and the migration of Th2-skewing CD301b^+^ dendritic cells (DCs) to draining lymph nodes (dLNs), initiating allergic immunity (Flayer et al., 2024; Perner et al., 2020; Serhan et al., 2019). These findings position sensory neurons as critical sentinels in allergen detection. Yet, in both natural and experimental settings, allergens are typically encountered via recurrent low-dose exposures (Custovic, 2015; Galli et al., 2008; Murrison et al., 2019; Ogulur et al., 2025). How these individually subthreshold encounters cumulatively drive allergic sensitization remains unclear.

Neurons are among the most energetically demanding cells in the body, relying heavily on mitochondrial oxidative phosphorylation (OXPHOS) to sustain excitability and neurotransmission (Kann and Kovacs, 2007; Trigo et al., 2022). Sensory neurons, particularly unmyelinated C-fibers, are especially vulnerable to energetic stress given their long axons and high ATP demands (Cheng et al., 2022; Wang et al., 2008). To meet these demands, sensory neurons employ adaptive metabolic strategies, including mitochondrial remodeling, acquisition, and biogenesis (Au et al., 2022; Guilford et al., 2017; van der Vlist et al., 2022; Van Laar et al., 2018; Xie et al., 2025). However, such adaptations can become maladaptive; in models of inflammatory pain, enhanced mitochondrial function drives the persistent pain sensitization phenomenon of hyperalgesic priming (Willemen et al., 2023). Whether similar metabolic priming occurs in response to allergen-induced itch remains unknown.

The mechanistic target of rapamycin complex 1 (mTORC1) is a central regulator of cellular metabolism that integrates diverse environmental inputs to coordinate mitochondrial function, protein synthesis, and energy production (Liu and Sabatini, 2020; Panwar et al., 2023). One key mechanism involves activation of the transcriptional coactivator peroxisome proliferator-activated receptor gamma coactivator-1α (PGC-1α), a master regulator of mitochondrial biogenesis. Through upregulating PGC-1α expression and activity, mTORC1 signaling drives mitochondrial biogenesis and enhances spare respiratory capacity, thereby enabling cells to meet increased energy demands (Cunningham et al., 2007; Marchetti et al., 2020). In neurons, mTORC1 supports axonal integrity and plasticity (Lipton and Sahin, 2014; Panwar *et al*., 2023; Querfurth and Lee, 2021), while aberrant activation contributes to chronic pain (Chen et al., 2022; Megat et al., 2019; Melemedjian et al., 2011; Xu et al., 2014). Given the high metabolic demands of sensory neurons and the role of mTORC1 in mitochondrial homeostasis and neuronal plasticity, mTORC1 is well-positioned to regulate how sensory neurons adapt to repeated environmental stimuli such as allergens.

Emerging evidence suggests that “trained immunity”, the long-term functional reprogramming of innate immune cells to enhance future responsiveness, extends beyond traditional immune cells to include non-immune cell types (Fanucchi et al., 2021; Lee et al., 2024; Netea et al., 2020; Ochando et al., 2023). We reasoned that a similar principle might apply to sensory neurons, which directly detect protease allergens and exhibit activity-dependent plasticity influenced by their metabolic state (Jaras et al., 2025). We therefore hypothesized that recurrent low-dose exposure to protease allergens would induce a form of innate neuroimmune “memory” in sensory neurons. In this model, initial allergen exposure would lead to sustained mTORC1 signaling, driving mitochondrial adaptations and neuronal hyper-responsiveness, thereby amplifying allergic immune initiation upon re-exposure. Consistent with this hypothesis, we found that repeated exposure to the model protease allergen papain established a neuron-intrinsic state of innate neuroimmune memory, marked by augmented itch, enhanced migration of Th2-skewing CD301b^+^ DC migration to dLNs, and increased Th2 differentiation. This innate memory state arose independently of adaptive immune cells or systemic cues but required initial activation of sensory neurons. Mechanistically, protease allergens induced sustained mTORC1 activation in sensory neurons, which, through PGC-1α-associated mitochondrial remodeling, conferred heightened responsiveness upon allergen re-exposure. Disruption of either mTORC1 signaling or mitochondrial stability abrogated training without affecting primary responses. Notably, neuroimmune training conferred heightened responses not only to the same allergen but also to antigenically distinct protease allergens. Together, these findings establish mTORC1-driven metabolic reprogramming in sensory neurons as a novel form of innate memory, providing a mechanistic basis for allergen cross-sensitization and the progressive accumulation of allergen sensitivities observed in atopic disease.

## RESULTS

### Repeated allergen exposure establishes a state of innate neuroimmune memory

To test whether allergens can induce a form of innate neuroimmune memory, we developed a repeated allergen exposure model adapted from protocols of hyperalgesic priming (Parada et al., 2003; Zhang et al., 2025). Wild-type C57BL/6 mice were intradermally (i.d.) injected with allergens at a 7-day interval. This time point was chosen to allow any acute effects of the first injection to resolve, thereby minimizing the confounding effects of acute inflammation at the injection site. Sensory neuronal activation was assessed by quantifying itch/scratch behavior in response to the model protease allergen papain, and immune responses were evaluated by DC migration and Th2 differentiation (**Figure 1A**). As expected, a single injection of papain (Pap), but not the inert protein ovalbumin (Ova), elicited acute scratching (Perner *et al*., 2020).

**Figure 1:**
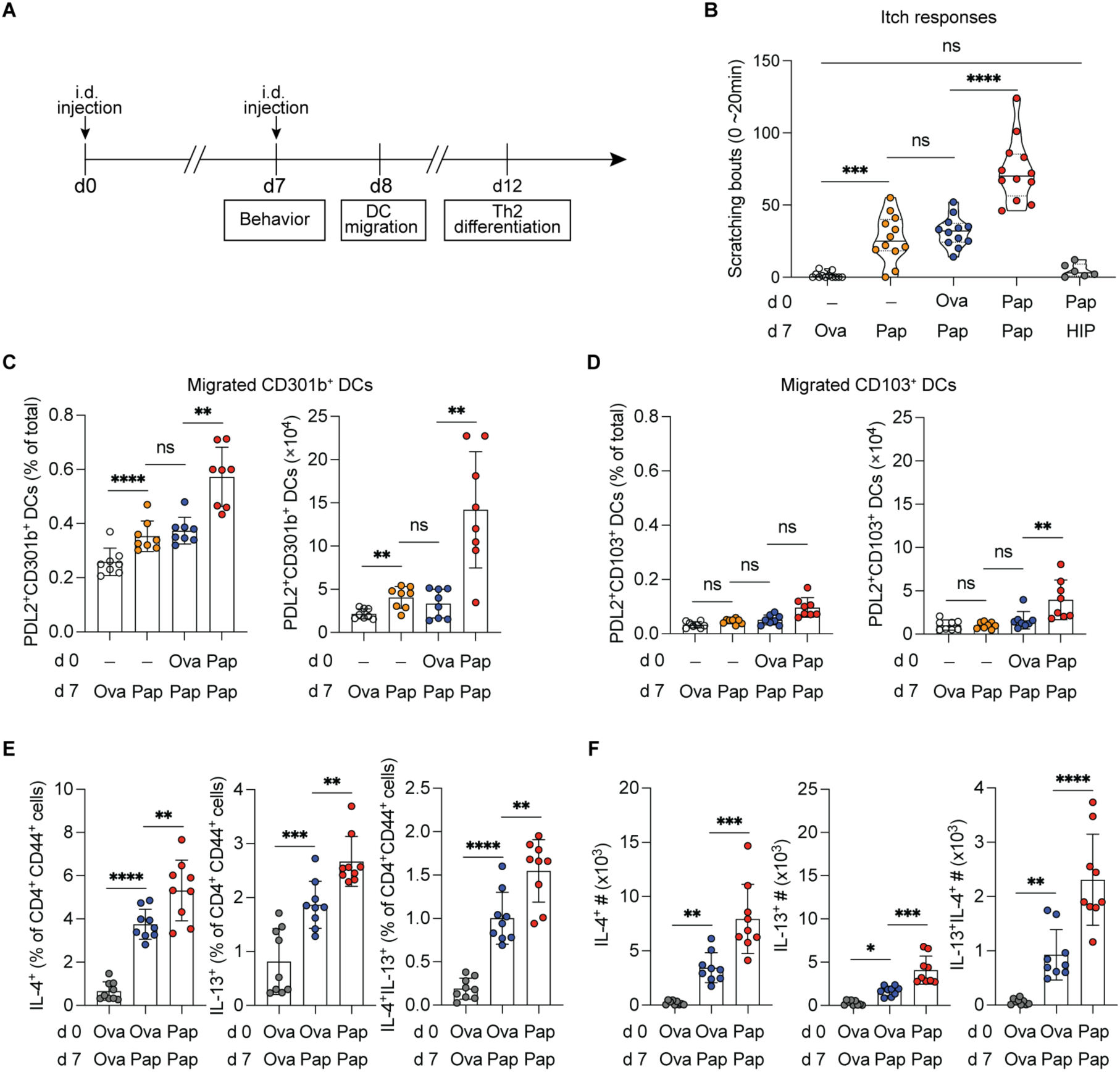
Repeated allergen exposure leads to neuroimmune memory. (**A**) Schematic of the allergen exposure protocol used to assess behavior, dendritic cell (DC) migration, and Th2 differentiation. Briefly, wild-type (WT) mice received intradermal (i.d.) injections as indicated into the ipsilateral cheek on days 0 and 7. Behavioral responses were recorded for 20 minutes immediately following injections as indicated on day 7. To assess DC migration, mice received i.d. footpad injections as indicated on days 0 and 7, and DC migration into the popliteal draining lymph node (dLN) was analyzed 22-24 hours after the secondary injections. Th2 differentiation in the popliteal dLN was evaluated on day 12, corresponding to 5 days after the secondary injections indicated on day 7. (**B**) WT mice received i.d. cheek injections with ovalbumin (Ova) or no injection (“–”) on day 0, followed by Ova, Papain (Pap), or heat-inactivated papain (HIP) on day 7. Scratching behavior was quantified as the total number of scratch bouts at the site of secondary injection. (**C-D**) Flow cytometric analysis of percentage and cell number of PDL2⁺CD301b⁺ and PDL2⁺CD103⁺ CD11c^+^ DCs in the popliteal dLN 22-24 hours after secondary injections as indicated. (**E-F**) Flow cytometric analysis of IL-4⁺, IL-13⁺, and IL-4⁺IL-13⁺ cells within the CD4⁺CD44^high^ T cell population in the popliteal dLN 5 days after secondary injections as indicated, shown as percentage (**E**) and cell number (**F**). Each data point represents an individual mouse. Violin plots show the median and quartiles. Bar plots are mean ± S.D. All data represent at least two independent experiments with each experiment including ≥3 mice per group. All plots show data combined from multiple experiments. Statistical tests: ordinary one-way ANOVA with Tukey’s multiple comparisons test (**B-F**). \*\**p*<0.01; \*\*\**p*<0.001; \*\*\*\**p*<0.0001; ns, not significant.

The itch response to papain given after an initial Ova injection was indistinguishable from that of a single papain injection, confirming that neither a non-protease antigen (Ova) nor the injection itself primes neuronal responses (**Figure 1B**). In contrast, mice exposed to papain on both day 0 and day 7 exhibited markedly enhanced scratching upon re-exposure (**Figure 1B**). This amplification was dependent on the protease activity of papain, as mice initially injected with papain but re-exposed to heat-inactivated papain (HIP) exhibited no detectable scratch response (**Figure 1B**). Thus, initial exposure to a proteolytically active allergen primes sensory neurons for heightened responsiveness upon re-exposure to the active allergen.

We next asked whether this neuronal training influences the initiation of allergic immunity. In the skin, CD301b^+^ DCs are known to drive Th2 responses, whereas CD103^+^ DCs include the XCR1^+^ DC subset that drives Th1 responses (Gao et al., 2013; Kumamoto et al., 2013; Lanca et al., 2022; Tussiwand et al., 2015). Prior studies have shown that allergen-activated sensory neurons, via local SP release, selectively promote the migration of activated, or PDL2^+^, Th2-skewing CD301b^+^ dermal DCs, but not CD103^+^ or XCR1^+^ DCs, from the skin to the dLNs (Flayer *et al*., 2024; Perner *et al*., 2020). To determine whether the enhanced sensory neuronal responses on recurrent allergen exposure similarly augmented DC responses, we performed flow cytometric analysis of Th1-skewing and Th2-skewing DCs in the popliteal dLN following repeated allergen exposure (**Supplementary Figure 1A**). Consistent with prior work, a single papain injection triggered the migration of activated CD301b^+^ DCs, but not CD103^+^ DCs (**Figure 1C-D**) (Flayer *et al*., 2024; Perner *et al*., 2020). However, repeated papain exposure further enhanced CD301b^+^ DC migration, while CD103^+^ DCs were minimally affected, indicating the biased amplification of Th2-skewing DC trafficking (**Figure 1C-D**). Because CD301b^+^ DC migration is essential for Th2 priming (Kumamoto *et al*., 2013; Sokol et al., 2018), we next examined T cell responses by flow cytometry (**Supplementary Figure 1B**). Repeated papain exposure significantly increased IL-4^+^ and IL-13^+^ CD4^+^ T cells compared to Ova-injected controls (**Figure 1E-F**). This effect was largely lost when re-exposure was performed with HIP (**Supplementary Figure 1C-D**), demonstrating that protease activity, but not antigenic sequence, was required. Together, these findings show that repeated protease allergen exposure establishes a state of neuroimmune memory, in which protease allergens promote enhanced itch, CD301b^+^ DC migration, and augmented Th2 differentiation upon allergen re-exposure. Having established the existence of an allergen-induced memory state, we next sought to determine whether this phenomenon is intrinsic to neurons or involves systemic or adaptive immune factors.

### Allergen-induced neuroimmune training is independent of adaptive lymphocytes

In the context of allergic inflammation, sensory neurons and immune cells engage in reciprocal communication through the exchange of soluble factors. Epithelial- and immune-derived mediators such as TSLP, LTC_4_, and IL-31 directly trigger itch, while IL-3, IL-4, and IL-13 prime sensory neuron responsiveness (Cevikbas et al., 2014; Flayer *et al*., 2024; Oetjen et al., 2017; Voisin et al., 2021; Wang et al., 2021; Wilson et al., 2013). To assess whether soluble or systemic factors contribute to the enhanced itch response observed upon allergen re-exposure, we examined whether neuroimmune training was site-specific. Mice injected on day 0 with papain into the right cheek exhibited robust scratching upon ipsilateral re-exposure on day 7 (**Figure 2A**). However, contralateral re-exposure failed to augment scratching (**Figure 2B**), indicating that the trained state is locally imprinted in the initially exposed skin site, rather than due to circulating factors.

**Figure 2:**
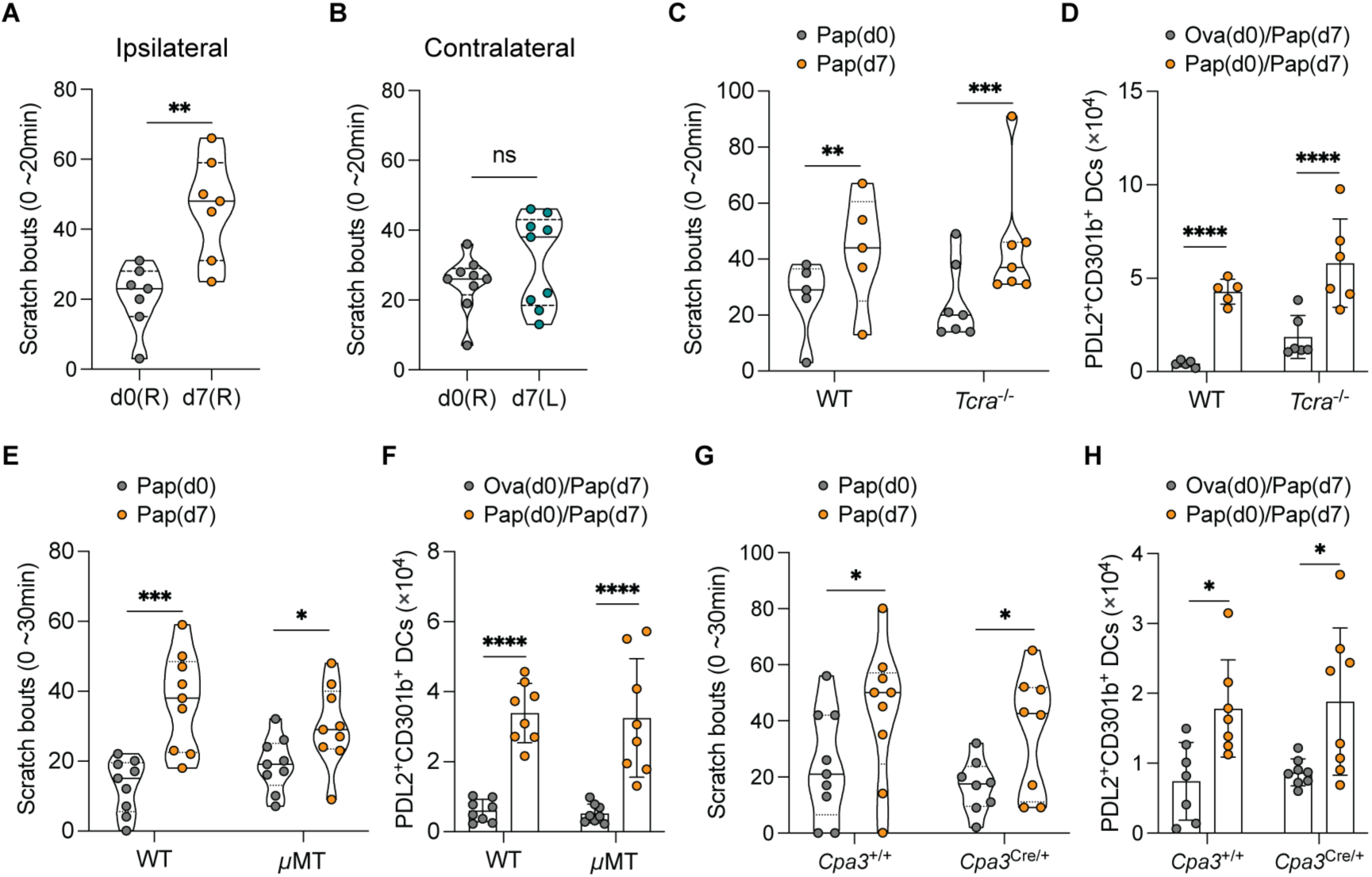
Allergen-induced neuroimmune memory occurs independently of lymphocytes, mast cells, or basophils. (**A-B**) WT mice were i.d. injected with papain (Pap) into one cheek on day 0, followed by ipsilateral or contralateral papain injection on day 7. Scratching behavior was quantified as the total number of scratch bouts at the site of secondary injection. (**C-H**) WT or *Tcra*^-/-^ mice (**C-D**), WT or *μ*MT^-/-^ mice (**E-F**), and littermate control *Cpa3*^+/+^ or *Cpa3*^Cre/+^ mice (**G-H**) received i.d. cheek injections as indicated on days 0 and 7. Scratching behavior was quantified as in (**A-B**). The number of PDL2⁺CD301b⁺ DCs in the popliteal dLN 22-24 hours after secondary footpad injections was analyzed by flow cytometry. Each data point represents an individual mouse. Violin plots show the median and quartiles. Bar plots are mean ± S.D. All data represent at least two independent experiments with each experiment including ≥3 mice per group. All plots show data combined from multiple experiments, except in (**C-D**) where one representative experiment is shown. Statistical tests: two-tailed unpaired *t*-test (**A-B**); two-way ANOVA with Tukey’s multiple comparisons test (**C-H**). \**p*<0.05; \*\**p*<0.01; \*\*\**p*<0.001; \*\*\*\**p*<0.0001; ns, not significant.

We next examined the role of adaptive lymphocytes. Both *Tcrα*^-/-^ mice (lacking αβ T cells) and *μMT*^-/-^ mice (lacking B cells) maintained enhanced itch responses and CD301b^+^ DC migration upon allergen re-exposure (**Figure 2C-F**). *Rag2^−/−^* mice (lacking all T and B cells) are known to have an impaired primary response to protease allergens because they lack IL-3-producing γδ T cells that normally help prime initial neuronal responses to allergens (Flayer *et al*., 2024). As expected, *Rag2^-/-^*mice exhibited a blunted initial response to allergen; however, they still exhibited enhanced scratch and DC migration upon rechallenge, indicating that the training mechanism does not require adaptive immunity (**Supplementary Figure 2A-B**). Finally, since mast cells and basophils mediate itch through both IgE- and Mas-related G protein-coupled receptor (MRGPR)-dependent pathways (Meixiong et al., 2019; Wang *et al*., 2021), we assessed training in *Cpa3*^Cre/+^ mice, which lack both cell types (Lilla et al., 2011). In these mice, repeated papain exposure still led to enhanced scratching and CD301b^+^ DC migration (**Figure 2G-H**). Together, these data show that allergen-induced neuroimmune training is independent of adaptive lymphocytes, mast cells, basophils, and systemic mediators, and suggest a non-immune mechanism.

### Allergen-induced neuroimmune training requires neuronal activation

Protease allergen-sensing neurons are enriched in their expression of the calcium channel transient receptor potential vanilloid subtype 1 (TRPV1) and the sodium channel Nav1.8 (encoded by *Scn10a*) (Flayer *et al*., 2024; Perner *et al*., 2020). Nav1.8 is expressed exclusively in sensory neurons, where it mediates depolarization and transmission of sensory stimuli, such as those detected by TRPV1. To determine whether sensory neuronal depolarization is required for the establishment of neuroimmune memory, we transiently blocked neuronal excitability during the initial allergen exposure using QX314, a lidocaine derivative that enters TRPV1- and TRPA1-expressing neurons and blocks sodium channel activation (Binshtok et al., 2007; Roberson et al., 2013). Mice injected with papain alone during the first exposure developed robust scratching upon re-exposure, whereas initial co-injection of QX314 completely abolished this augmentation, returning responses to baseline levels (**Figure 3A**). Using Kaede transgenic mice to track migration of photoconverted (Kaede^Red^) skin DCs (Tomura et al., 2008), we found that neuronal blockade with QX314 during the initial allergen exposure similarly abrogated the enhanced migration of CD301b⁺ DCs upon allergen re-exposure (**Figure 3B**). Similar results were obtained with A-803467, a selective Nav1.8 sodium channel blocker (Jarvis et al., 2007), which abrogated both enhanced scratching and CD301b⁺ DC migration upon re-exposure (**Figure 3C-D**). Thus, neuronal activation at the time of the first allergen exposure is essential for the establishment of allergen-induced neuroimmune memory.

**Figure 3:**
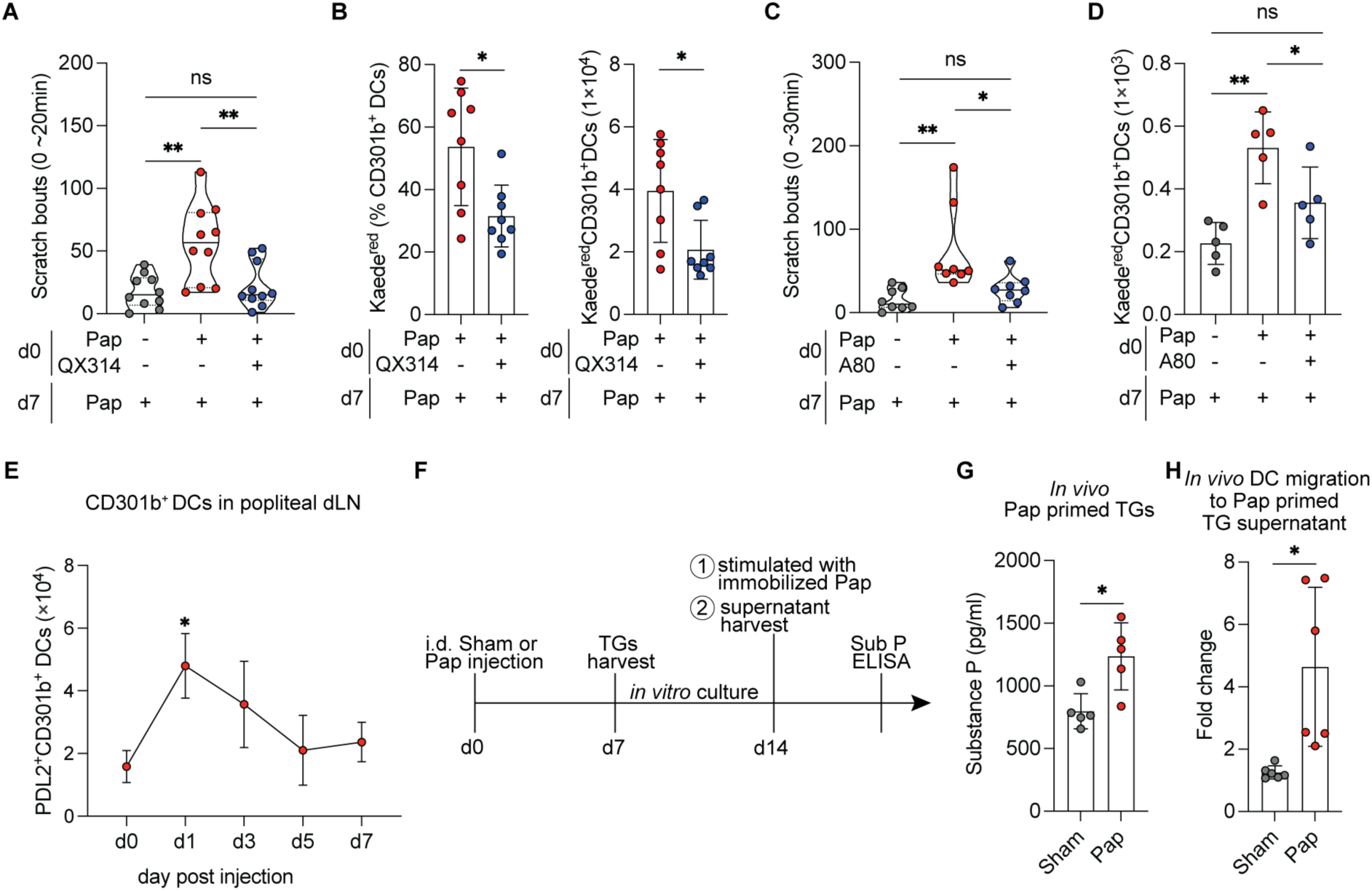
Allergen-induced neuroimmune memory requires initial neuronal activation. (**A**) WT mice received i.d. cheek injections of papain (Pap) or papain co-administered with QX314 on day 0, followed by a secondary papain injection on day 7. Scratching behavior was quantified as the total number of scratch bouts at the site of secondary injection. (**B**) Kaede^GFP^ mice received i.d. footpad injections of papain or papain co-administered with QX314 on day 0 and then were photoconverted at the initial injection site followed by papain injection on day 7. The percentage and total number of Kaede^Red^CD301b⁺ DCs in the popliteal dLN were quantified by flow cytometry. (**C**) WT mice were intraperitoneally (i.p.) injected with vehicle or A-803467 (A80) 4 hours prior to indicated i.d. cheek injections on day 0, followed by papain injections on day 7. Scratching bouts were quantified as described in (**A**). (**D**) Kaede^GFP^ mice were i.p. injected with vehicle or A-803467 (A80) 4 hours prior to indicated i.d. footpad injections on day 0 and then were photoconverted at the initial injection site followed by papain injection on day 7. The total number of Kaede^Red^CD301b⁺ DCs in the popliteal dLN was quantified by flow cytometry. (**E**) Flow cytometric analysis of PDL2^+^CD301b⁺ DCs in the popliteal dLN at indicated time points following a single i.d. footpad injection of papain on day 0. (**F**) Schematic illustrating the stimulation of *in vitro*-cultured trigeminal ganglion (TG) neurons and the collection of cell-free supernatant for Substance P (Sub P) ELISA and subsequent injections in (G-H). (**G**) Substance P levels in the supernatant from sham (Ova) or papain-primed TG neurons stimulated *in vitro* with papain, measured by ELISA. (**H**) Fold change in numbers of Kaede^Red^CD301b⁺ DCs per dLN quantified by flow cytometry after migration from photoconverted skin of Kaede^GFP^ mice injected with cell-free supernatant derived from either sham or papain-primed TG neurons as shown in (**F**). Fold change was calculated by dividing dLN cell counts of indicated experimental condition over dLN cell counts obtained after footpad injection with media from cell-free culture wells. Each data point represents an individual mouse. Violin plots show the median and quartiles. Bar plots are mean ± S.D. All data represent at least two independent experiments with each experiment including ≥3 mice per group. All plots show data combined from multiple experiments, except in (**D-E**) and (**G-H**) where one representative experiment is shown. Statistical tests: ordinary one-way ANOVA with Tukey’s multiple comparisons test (**A, C, D, E**); two-tailed paired *t*-test (**B**); two-tailed unpaired *t*-test (**G, H**). \**p*<0.05; \*\**p*<0.01; ns, not significant.

We next asked how repeated allergen exposure amplifies CD301b⁺ DC migration. Flow cytometric analysis showed that CD301b⁺ DC numbers in the dLNs returned to baseline within days after the initial allergen exposure (**Figure 3E**), ruling out DC retention as a cause of increased accumulation in the lymph node. Instead, trigeminal ganglia (TG) neurons from papain-injected mice released significantly higher levels of SP when restimulated *in vitro* with immobilized papain compared to naïve controls (**Figure 3F-G**), suggesting that prior allergen exposure enhances neuropeptide release upon re-exposure. To test whether these neuronal factors are sufficient to drive augmented DC migration, we applied supernatants from papain-restimulated TG cultures to the photoconverted skin of Kaede mice. Supernatants from TG neurons previously exposed to papain induced significantly greater CD301b^+^ DC migration to dLNs than supernatants from sham-exposed TG neurons (**Figure 3H**). Together, these findings show that allergen-induced neuroimmune training requires initial sensory neuronal activation, which primes neurons for enhanced SP release upon re-exposure and thereby amplifies Th2-skewing DC migration and downstream allergic immune responses.

### Protease allergen stimulation induces sustained mTORC1 activation in sensory neurons

To define mechanisms underlying allergen-induced neuroimmune training in sensory neurons, we performed bulk RNA sequencing on pooled TGs from wild-type mice 7 days after intradermal exposure to Ova (sham), papain (Pap), or papain plus QX314 (Pap/QX314). Bulk RNA sequencing revealed minimal overall transcriptional differences at this time point (**Table 1-2, Supplementary Figure 3A-B**). However, gene set enrichment analysis (GSEA) identified the upregulation of metabolic pathways, including fatty acid metabolism and mTORC1 signaling, in the papain-exposed group compared to controls (**Figure 4A, Supplementary Figure 3C**). Notably, papain co-administration with QX314 attenuated these signatures (**Figure 4A**), indicating that neuronal activation is required for mTORC1 pathway induction.

**Table 1.**
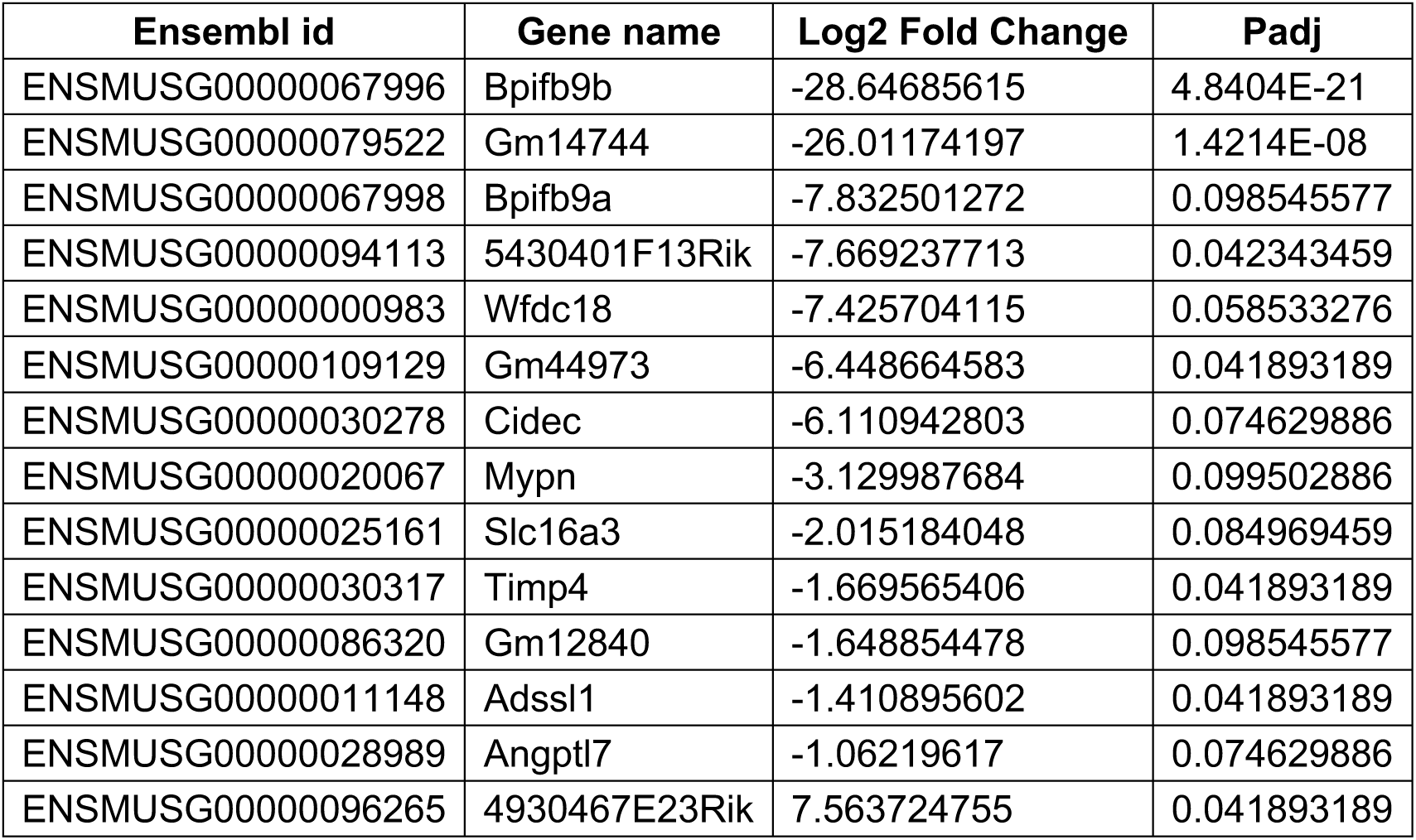
Genes with significant changes in the Pap group compared to the sham group.

**Table 2.**
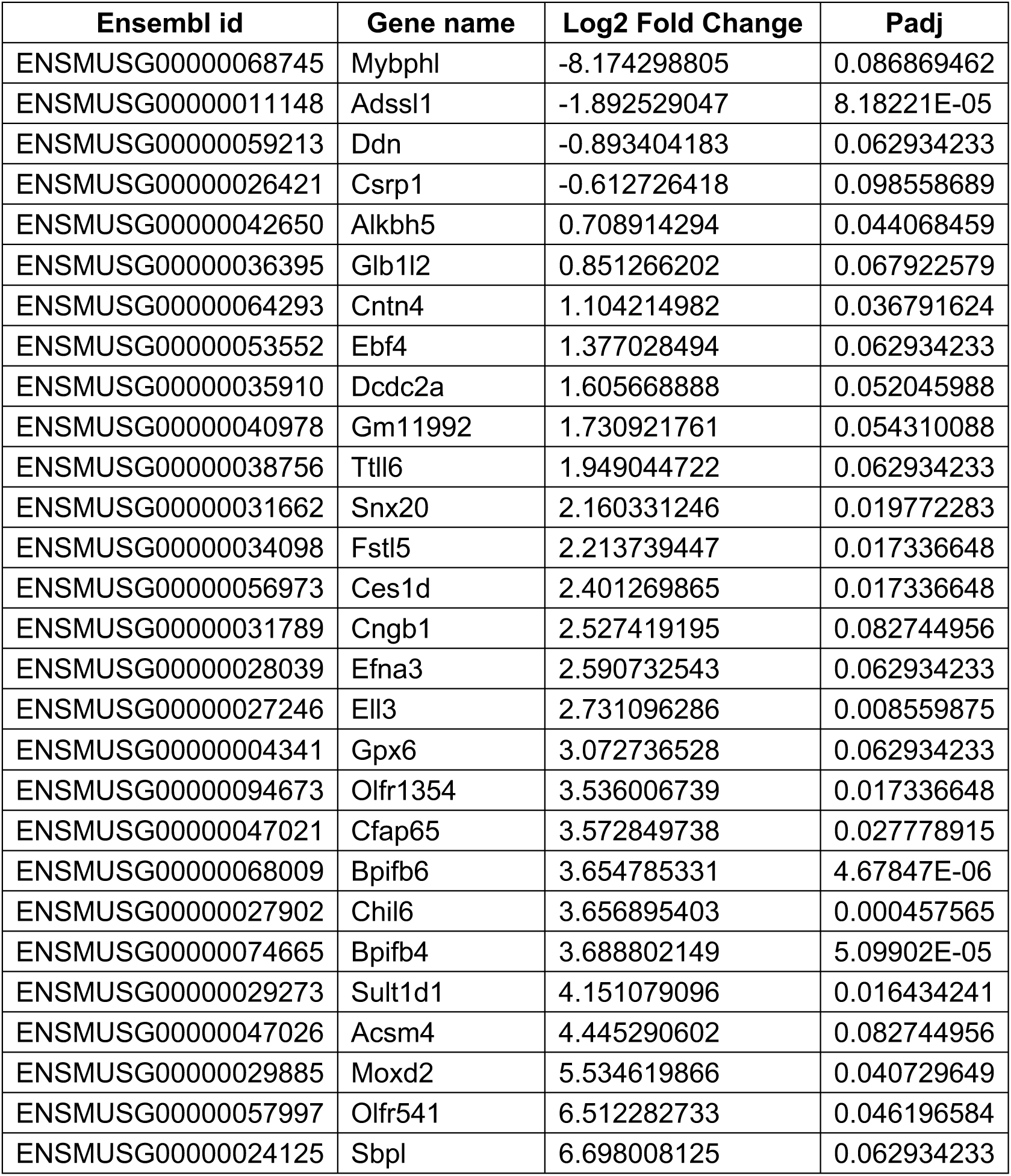
Genes with significant changes in the Pap group compared to the Pap/QX314 group.

**Figure 4:**
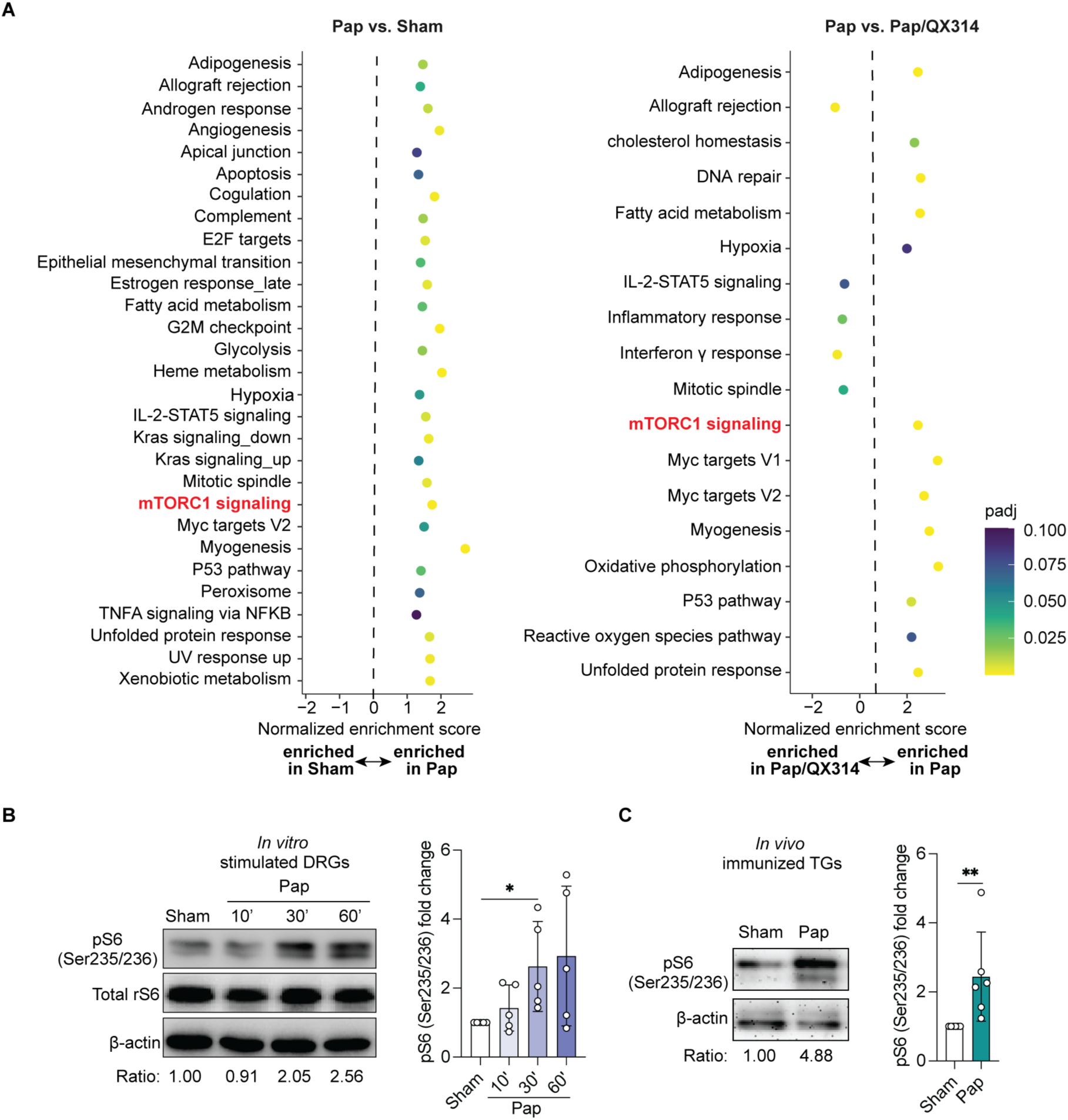
Protease allergen-stimulated sensory neurons exhibit mTORC1 activation. (**A**) Gene set enrichment analysis (GSEA) of hallmark signaling pathways in TG neurons 7 days after a single i.d. injection of ovalbumin (Sham), papain (Pap), or papain co-administered with QX314 (Pap/QX314) into the cheek. Bubble plot depicts normalized enrichment scores and significance levels. The left panel shows genes enriched in the Pap vs. Sham group, and the right panel shows genes enriched in the Pap vs. Pap/QX314 group. (**B-C**) Representative immunoblots (left) and quantification (right) of pS6 (Ser235/236) levels in DRG neurons stimulated *in vitro* with papain for the indicated time (**B**) and in TG neurons isolated from WT mice 7 days after a single i.d. papain cheek injection (**C**). The normalized pS6/β-actin ratio is shown below each lane. Bulk-seq in (**A**) was performed with three replicates per group. Each dot in the bar plot of (**B-C**) represents one independent assay. Bar plots are mean ± S.D. Statistical analysis: two-tailed ratio paired *t*-test (**B**, **C**); \**p*<0.05; \*\**p*<0.01.

mTORC1 is a master regulator of cellular metabolism whose activity is commonly monitored by phosphorylation of ribosomal protein S6 (pS6) (Liu and Sabatini, 2020; Panwar *et al*., 2023). Guided by our transcriptomic results, we examined pS6 expression in primary dorsal root ganglia (DRG) neurons stimulated *in vitro* with papain. Papain induced robust pS6 expression within 30 minutes after treatment (**Figure 4B**), which was abolished by co-treatment with the mTORC1 inhibitor sirolimus (also known as rapamycin) (**Supplementary Figure 3D**), confirming that this papain-induced S6 phosphorylation is mTORC1-dependent. *In vivo* cheek administration of papain led to an approximately two-fold increase in pS6 expression that persisted in TG neurons 7 days after papain exposure, demonstrating sustained mTORC1 activation (**Figure 4C**). Together, these data demonstrate that protease allergen stimulation induces prolonged mTORC1 signaling in sensory neurons, providing a potential mechanistic link between initial neuronal activation and the establishment of neuroimmune memory.

### Sensory neuronal mTORC1 activation is required for allergen-induced neuroimmune memory

Previous studies have shown that mTORC1 supports neuronal plasticity (Lipton and Sahin, 2014; Panwar *et al*., 2023), which can have pathologic outcomes in chronic pain by promoting neuronal responses to non-painful stimuli (Chen *et al*., 2022; Megat *et al*., 2019; Melemedjian *et al*., 2011; Xu *et al*., 2014). Based on these observations, we hypothesized that sustained mTORC1 activation in sensory neurons mediates allergen-induced neuroimmune memory. To test this, we first employed two structurally distinct mTORC1-specific inhibitors, sirolimus (also known as rapamycin) and temsirolimus (also known as CCI-779 or Torisel). As opposed to a previous report examining itch responses to chemical pruritogens (Obara et al., 2015), acute papain-induced scratching and CD301b^+^ DC migration on day 0 were unaffected by sirolimus or temsirolimus (**Figure 5A-B; Supplementary Figure 4A-B**). However, both drugs abrogated the enhanced scratching and augmented CD301b⁺ DC migration seen upon papain re-exposure on day 7 (**Figure 5A-B; Supplementary Figure 4B**). Thus, mTORC1 is dispensable for acute responses but required for the primed memory state.

**Figure 5:**
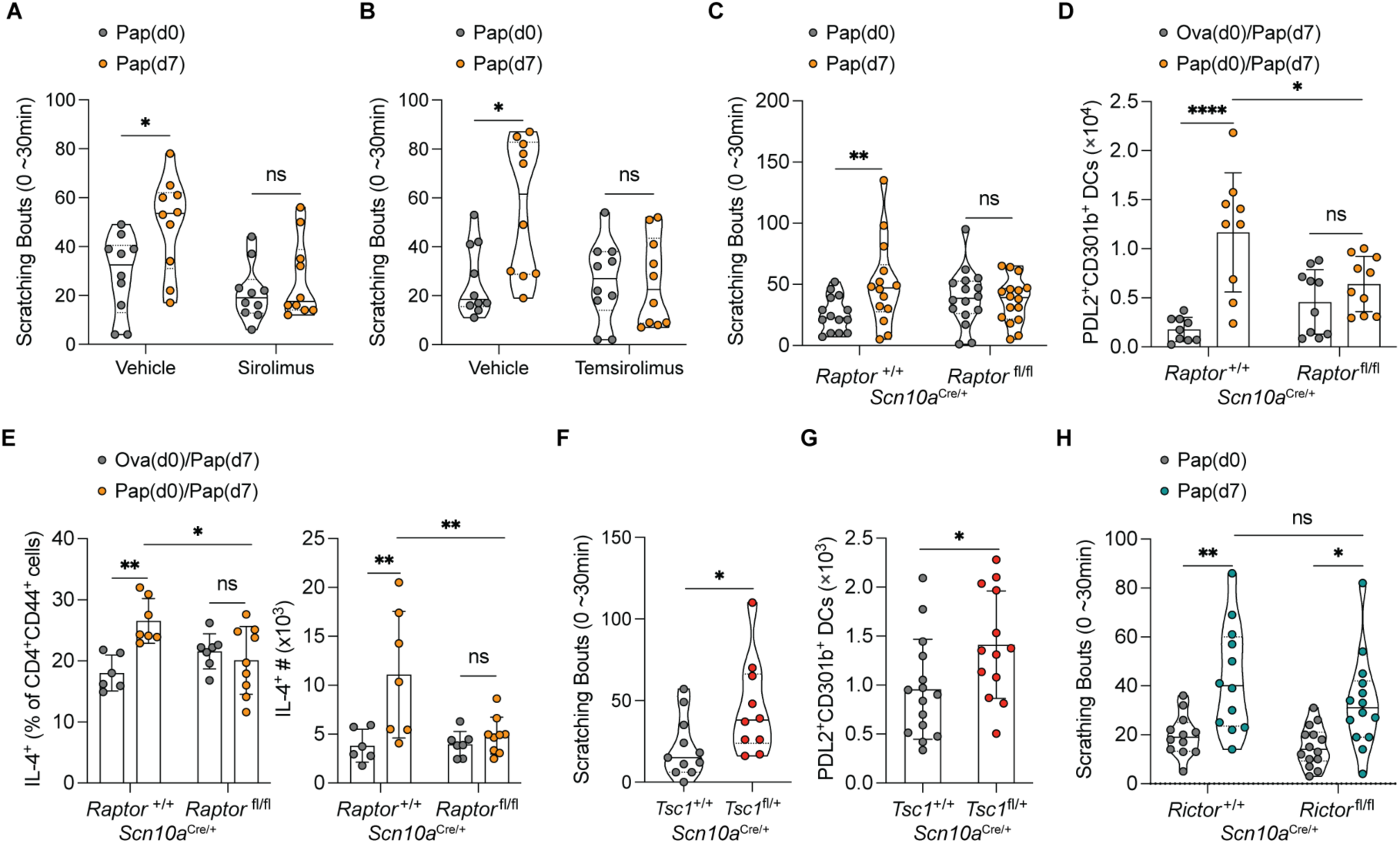
Sensory neuronal mTORC1 activation is required for allergen-induced neuroimmune memory. (**A-B**) WT mice received i.d. cheek injections of sirolimus, temsirolimus, or vehicle 2 hours before papain (Pap) cheek injections on day 0, followed by daily dosing on days 1 through 6 and a final dose 2 hours before secondary papain cheek injections on day 7. Scratching behavior was quantified as the total number of scratch bouts at the site of secondary injection. (**C-D**) Scratching responses (**C**) and flow cytometric analysis of PDL2⁺CD301b⁺ DCs in the popliteal dLN (**D**) from littermate control *Scn10a*^Cre/+^*Raptor*^+/+^ and *Scn10a*^Cre/+^*Raptor*^fl/fl^ mice after indicated injections on days 0 and 7. Scratching bouts were quantified as in (**A**). (**E**) Percentage and cell number of IL-4⁺ cells within the CD4⁺CD44^high^ T cell population in the popliteal dLN, as assessed by flow cytometry following footpad injections as indicated on days 0 and 7. (**F**) Scratching bouts were quantified as in (**A**) following a single papain cheek injection in *Scn10a*^Cre/+^*Tsc1*^fl/*+*^ mice and littermate control *Scn10a*^Cre/+^*Tsc1*^+/*+*^ mice. (**G**) Flow cytometric quantification of PDL2^+^CD301b⁺ DCs in the popliteal dLN after a single papain footpad injection in *Scn10a*^Cre/+^*Tsc1*^fl/*+*^ mice and littermate control *Scn10a*^Cre/+^*Tsc1*^+/*+*^ mice. (**H**) Scratching bouts were quantified as in (**A**) for *Scn10a*^Cre/+^*Rictor*^fl/fl^ mice and littermate control *Scn10a*^Cre/+^*Rictor*^+/+^ mice following cheek injections as indicated on days 0 and 7. Each data point represents an individual mouse. Violin plots show the median and quartiles. Bar plots are mean ± S.D. All data represent at least two independent experiments with each experiment including ≥3 mice per group. All plots show data combined from multiple experiments. Statistical tests: two-way ANOVA with Tukey’s multiple comparisons test (**A-E, H**) and two-tailed unpaired *t*-test (**F, G**). \**p*<0.05; \*\**p*<0.01; \*\*\*\**p*<0.0001; ns, not significant.

To confirm these findings genetically, we deleted *Raptor*, an essential component of the mTORC1 complex (Kim et al., 2002), in Nav1.8^+^ sensory neurons (*Scn10a*^Cre/+^*Raptor*^fl/fl^). These mice exhibited normal itch, CD301b^+^ DC migration, and Th2 differentiation upon initial papain exposure and displayed intact cutaneous innervation compared to littermate controls (**Figure 5C-E, Supplementary Figure 4C-D**). In contrast, the augmented scratching, enhanced CD301b⁺ DC migration, and increased Th2 differentiation normally seen upon papain re-exposure were absent in *Scn10a*^Cre/+^*Raptor*^fl/fl^ mice (**Figure 5C-E**). Conversely, persistent sensory neuronal mTORC1 activation, achieved by heterozygous deletion of the mTORC1 inhibitor *Tsc1* in Nav1.8^+^ neurons (*Scn10a*^Cre/+^*Tsc1*^fl/+^ mice) (Yang et al., 2006), was sufficient to elicit enhanced scratching and CD301b⁺ DC migration after a single papain exposure (**Figure 5F-G**), phenocopying a wild-type trained response. Together, these results demonstrate that sensory neuron-intrinsic mTORC1 signaling is both necessary and sufficient to induce the allergen-induced neuroimmune memory state.

mTOR signaling is mediated through two functionally distinct complexes, mTORC1 and mTORC2 (mammalian target of rapamycin complex 2) (Szwed et al., 2021). *Rictor*, a defined and essential component of mTORC2, has also been implicated in neuronal development and function (Liu and Sabatini, 2020; Querfurth and Lee, 2021; Switon et al., 2017). Because prolonged sirolimus exposure can indirectly inhibit mTORC2 (Sarbassov et al., 2006), we next assessed whether mTORC2 contributes to training. Papain induced only a transient increase in phosphorylated Akt (Ser473), a canonical mTORC2 target (Sarbassov et al., 2005), in cultured DRG neurons (**Supplementary Figure 4E**). Moreover, conditional deletion of the essential mTORC2 component *Rictor* in Nav1.8^+^ neurons (*Scn10a*^Cre/+^*Rictor*^fl/fl^) had no effect on either acute or memory responses (**Figure 5H; Supplementary Figure 4F**). Collectively, these findings establish that mTORC1 – but not mTORC2 – activation in sensory neurons is required for allergen-induced neuroimmune memory. While mTORC1 is dispensable for acute itch and initial allergic responses, it is essential for the long-term neuronal adaptations that amplify allergic reactivity.

### mTORC1 promotes mitochondrial remodeling required for allergen-induced neuroimmune memory

To investigate how mTORC1 supports neuroimmune memory, we focused on the transcriptional coactivator PGC-1α, a master regulator of mitochondrial biogenesis and oxidative metabolism (Qian et al., 2024). Because mTORC1 directly regulates PGC-1α expression and activity, enabling appropriate metabolic adaptation in response to environmental and energetic stressors (Bennett et al., 2022), we asked whether allergen exposure induced this axis in sensory neurons. Consistent with sustained mTORC1 activation (**Figure 4C**), initial papain exposure significantly upregulated PGC-1α protein in TG neurons within 24 hours, and this increase persisted for at least 7 days (**Figure 6A; Supplementary Figure 5A**). Sirolimus treatment partially abrogated PGC-1α induction (**Figure 6B; Supplementary Figure 5B**), confirming its dependence on mTORC1 signaling. PGC-1α upregulation promotes mitochondrial biogenesis and network connectivity, features associated with enhanced oxidative metabolism (Cunningham *et al*., 2007; Marchetti *et al*., 2020; Qian *et al*., 2024). In line with this, papain cheek-injected TG neurons exhibited a significant increase in mitochondrial DNA (mtDNA) copy number compared to sham controls, indicating augmented mitochondrial replication (**Supplementary Figure 5C**). Immunostaining for the mitochondrial outer membrane marker translocase of outer membrane-20 (TOM20) further revealed papain cheek-injected TG neurons exhibited enlarged, elongated and more complex mitochondria, reflected by increased aspect ratio (measure of length), form factor (combined measure of length and degree of branching) and mean branch length compared to sham controls (**Figure 6C; Supplementary Figure 5D-E**) (Chaudhry et al., 2020; Koopman et al., 2005; Rumbeiha et al., 2023). Together, these results indicate that allergen exposure activates the mTORC1-PGC-1α axis to drive mitochondrial biogenesis and remodeling in sensory neurons.

**Figure 6:**
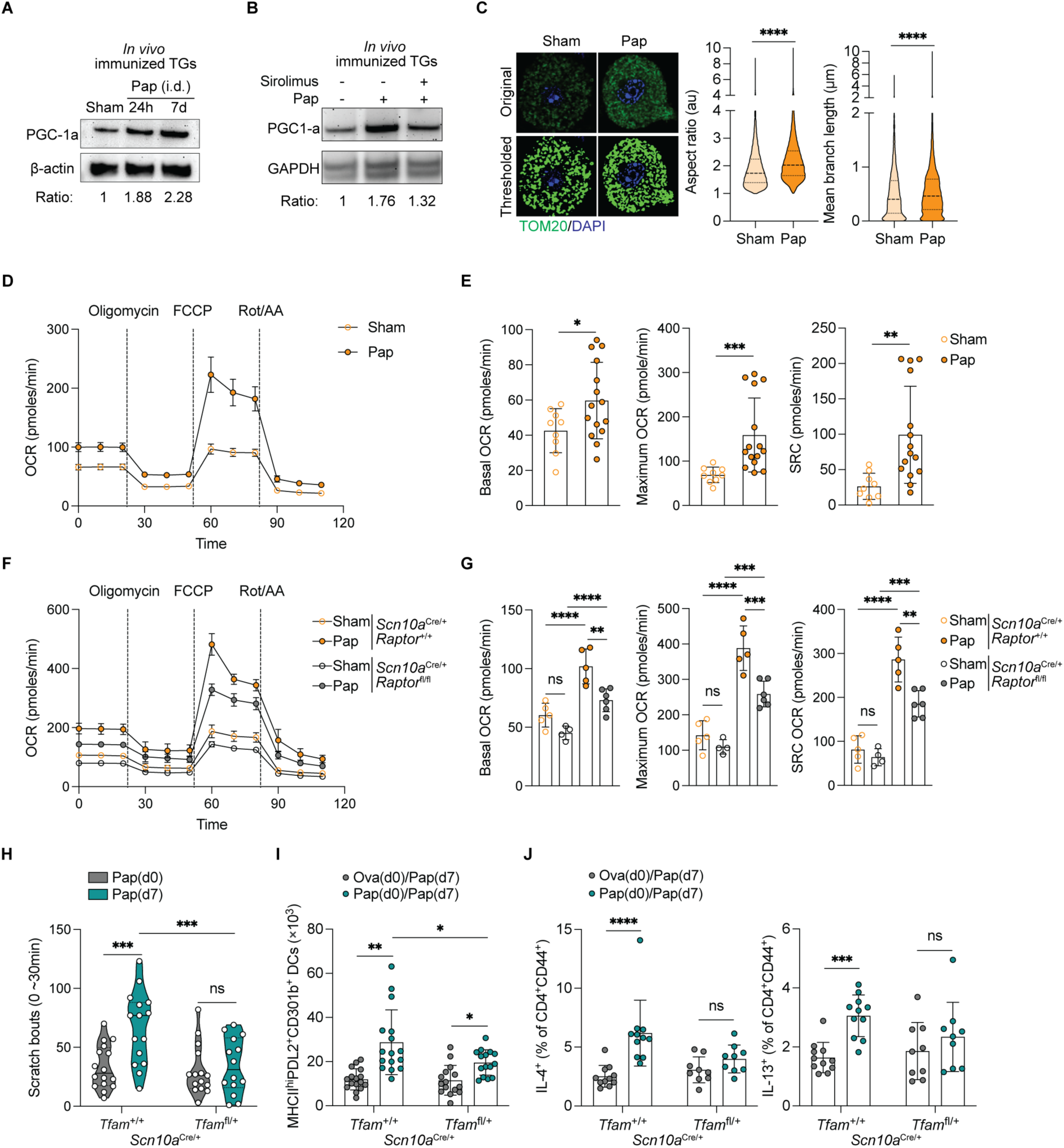
mTORC1 promotes mitochondrial function required for allergen-induced neuroimmune memory. (**A**) Representative immunoblots of PGC-1α in TG neurons from WT mice 24 hours (h) or 7 days (d) after i.d. cheek injection of ovalbumin (Sham) or papain (Pap). Normalized PGC-1α/β-actin ratios are shown under each lane. (**B**) Representative immunoblots of PGC-1α in TG neurons harvested 7 days after papain injection with or without sirolimus treatment. WT mice received i.d. cheek treatment with vehicle or sirolimus 2 hours before i.d. papain injection on day 0, followed by an additional i.d. cheek treatment on day 1 to ensure complete inhibition of early allergen-induced mTORC1 activation. Normalized PGC-1α/GAPDH ratios are shown under each lane. (**C**) Confocal images (left) of TOM20 in TG neurons isolated from WT mice 7 days after i.d. cheek injection of sham (Ova) or papain. The upper panels show original images, and the lower panels show thresholded images generated using the Mitochondrial Analyzer plugin in Image J. Violin plots (right) show quantification of mitochondrial aspect ratio (arbitrary units, au) and mean branch length (μm) in sham (170 neurons) or papain-primed (194 neurons) TG neurons obtained with the same plugin. (**D-E)** Oxygen consumption rate (OCR) of TG neurons isolated from WT mice after treatment regimen described in (**C**), measured by Seahorse metabolic flux analysis. Responses to oligomycin, FCCP (carbonyl cyanide-4-(trifluoromethoxy) phenylhydrazone), Rot/AA (rotenone and antimycin A) are shown (**D**) with corresponding statistical analysis (**E**). (**F-G**) OCR profiles of TG neurons isolated from *Scn10a*^Cre/+^*Raptor*^fl/fl^ and *Scn10a*^Cre/+^*Raptor*^+/+^ littermate control mice 7 days after i.d. cheek injection of sham or papain, assessed by Seahorse metabolic flux analysis. Responses to oligomycin, FCCP, and Rot/AA are shown (**F**), with corresponding statistical analysis (**G**). Sham groups for each genotype are indicated by open circles, and papain groups by filled circles. (**H-J**) Scratching responses (**H**), cell number of PDL2⁺CD301b⁺ DCs (**I**), and percentage of IL-4⁺ and IL-13⁺ cells within the CD4⁺CD44^high^ T cell compartment in the popliteal dLN (**J**) from *Scn10a*^Cre/+^*Tfam*^fl/*+*^ and *Scn10a*^Cre/+^*Tfam*^+/*+*^ littermate control mice following injections as indicated on days 0 and 7. Scratching behavior was quantified as the total number of scratch bouts at the site of secondary injection. Each data point represents an individual mouse. Violin plots show the median and quartiles. Bar plots are mean ± S.D. All data represent at least two independent experiments with each experiment including ≥3 mice per group. All plots show data combined from multiple experiments. Statistical tests: two-tailed unpaired *t*-test (**C, E**); ordinary one-way ANOVA with Tukey’s multiple comparisons test (**G**); two-way ANOVA with Tukey’s multiple comparisons test (**H-J**). \**p*<0.05; \*\**p*<0.01; \*\*\**p*<0.001; \*\*\*\**p*<0.0001; ns, not significant.

We next asked whether this remodeling enhances mitochondrial respiration. Recent studies have linked hyperalgesic priming to enhanced mitochondrial activity, characterized by increased spare respiratory capacity (SRC) and elevated mitochondrial NAD⁺/NADH ratios (Willemen *et al*., 2023). Seahorse analysis of TG neurons 7 days after i.d. cheek injection with papain revealed significantly increased basal and maximal oxygen consumption rates (OCR), ATP production, and SRC compared to Ova, or sham, injected controls (**Figure 6D-E, supplementary Figure 5F**). Glycolytic activity as measured by the extracellular acidification rate (ECAR) remained largely unchanged following papain stimulation (**Supplementary Figure 5G**), indicating that allergen-induced metabolic adaptation favors oxidative phosphorylation (OXPHOS) rather than glycolysis. Importantly, TG neurons from *Scn10a*^Cre/+^*Raptor*^fl/fl^ mice exhibited defective allergen-induced OXPHOS enhancement, despite maintaining baseline respiration and glycolysis (**Figure 6F-G; Supplementary Figure 5H-I**). Thus, mTORC1 drives allergen-induced mitochondrial remodeling and enhanced respiratory capacity. To test whether mitochondrial stability and responsiveness are required for training, we examined *Scn10a*^Cre/+^*Tfam*^fl/+^ mice. *Tfam* is a nuclear-encoded mitochondrial transcription factor required for mitochondrial DNA replication; haploinsufficiency leads to reduced mitochondrial content and stability (Koh et al., 2021; Kozhukhar and Alexeyev, 2023; Kuroda et al., 2021). *Scn10a*^Cre/+^*Tfam*^fl/+^ mice, in which *Tfam* haploinsufficiency is restricted to Nav1.8^+^ sensory neurons, exhibited normal acute itch and activated CD301b^+^ DC migration upon initial papain exposure, but failed to develop the enhanced scratching, augmented DC migration, or increased Th2 differentiation observed in controls upon re-exposure (**Figure 6H-J**). Together, these findings demonstrate that mTORC1-PGC-1α signaling promotes mitochondrial remodeling and enhanced oxidative metabolism in sensory neurons, and that mitochondrial stability in sensory neurons is required for the establishment of allergen-induced neuroimmune memory.

### Allergen-induced neuroimmune training enables antigen-independent cross-sensitization

Our findings suggest that mTORC1-mediated neuroimmune training, characterized by the augmented itch responses, enhanced CD301b^+^ DC migration, and increased Th2 differentiation, depends on the protease activity of allergens, rather than antigen recognition. We therefore hypothesized that structurally distinct allergens sharing cysteine protease activity could induce cross-sensitization of neuroimmune responses independent of antigen specificity. To test this, we compared papain, a plant-derived cysteine protease, and Der p 1, a house dust mite-derived cysteine protease, as antigenically unrelated, but enzymatically similar allergens (Soh *et al*., 2023). To mimic real-world, low-dose repeated exposure as well as to normalize between papain and house dust mite (HDM) protease activity, mice were initially injected with low-dose allergens (or Ova as sham) for three consecutive days (days 0-2), followed by HDM exposure on day 9 (**Figure 7A**). As expected, repeated HDM exposure induced neuroimmune memory, as evidenced by enhanced scratching, increased migration of activated CD301b^+^ DCs, and augmented Th2 differentiation compared to Ova (sham) controls (**Figure 7B-D**). Strikingly, despite no antigen overlap between papain and HDM, the papain-primed mice responded to HDM as if they had initially been exposed to HDM (**Figure 7B-D**). To confirm that the papain-primed HDM response was truly HDM-specific, we restimulated lymph node cultures with heat-inactivated HDM (HI-HDM) and found that HDM and papain priming both led to equivalent HDM-specific IL-4 and IL-13 production (**Figure 7E**). Thus, protease activity alone enables cross-sensitization between distinct allergens.

**Figure 7:**
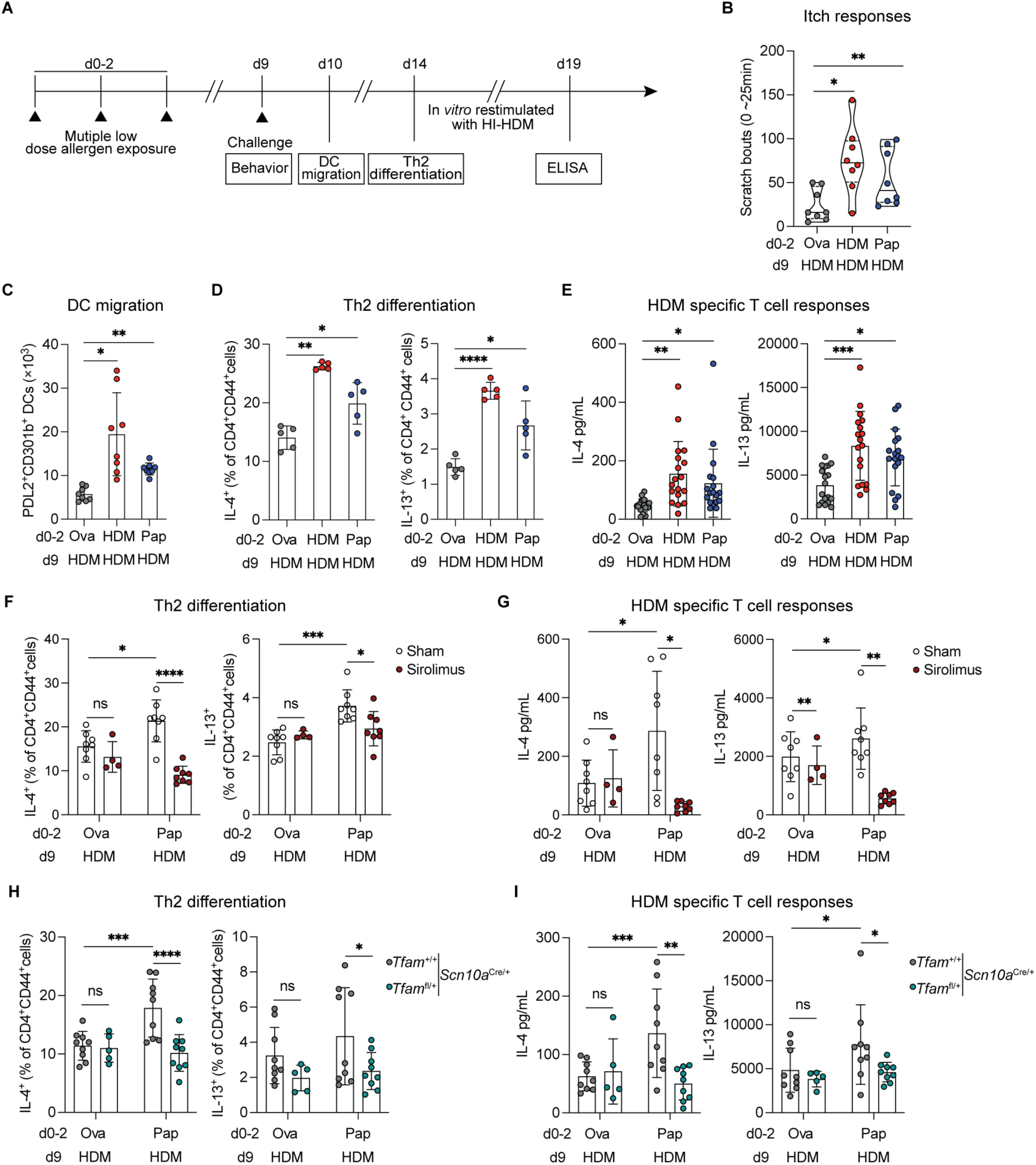
Allergen-induced neuroimmune memory promotes responsiveness to recurrent low dose allergen exposure in an antigen-independent manner. (**A**) Schematic of the recurrent low dose exposure model used to assess behavioral responses, DC migration, Th2 differentiation, and house dust mite (HDM)-specific cytokine production by ELISA. (**B**) Scratching behavior was quantified as the total number of ipsilateral cheek scratch bouts following HDM cheek injection on day 9. (**C**) Flow cytometric analysis of PDL2⁺CD301b⁺ DCs in the popliteal dLN of WT mice that received i.d. cheek injections as indicated on days 0-2 and 9. (**D**) Frequency and absolute number of IL-4⁺ and IL-13^+^ cells within the CD4⁺CD44^high^ T cell population in the popliteal dLN, as assessed by flow cytometry in WT mice that received i.d. footpad injections as indicated on days 0-2 and 9. (**E**) Whole popliteal dLN cells were harvested from WT mice after indicated footpad injections and were restimulated *in vitro* with heat-inactivated HDM (HI-HDM). HDM-specific IL-4 and IL-13 in the culture supernatant was quantified by ELISA. (**F**) Frequency and absolute number of IL-4⁺ and IL-13⁺ cells within the CD4⁺CD44^high^ T cell population in the popliteal dLN of WT mice following i.d. footpad injections as indicated on days 0-2 and 9 with or without sirolimus treatment. (**G**) ELISA quantification of HDM-specific IL-4 and IL-13 from popliteal dLN cultures as in (**E**), in WT mice that received injections as indicated. (**H**) Flow cytometric analysis of IL-4⁺ and IL-13⁺ CD4⁺CD44^high^ T cells in the popliteal dLN of *Scn10a*^Cre/+^*Tfam*^fl/*+*^ and *Scn10a*^Cre/+^*Tfam*^+/*+*^ littermate control mice after injections into the footpad as indicated. (**I**) ELISA quantification of HDM-specific IL-4 and IL-13 from popliteal dLN cultures as in (E) in *Scn10a*^Cre/+^*Tfam*^fl/*+*^ and *Scn10a*^Cre/+^*Tfam*^+/*+*^ littermate control mice after i.d. footpad injections as indicated. Each data point represents an individual mouse. Violin plots show the median and quartiles. Bar plots are mean ± S.D. All data represent at least two independent experiments with each experiment including ≥3 mice per group. All plots show data combined from multiple experiments, except in (**D**) where one representative experiment is shown. Statistical tests: ordinary one-way ANOVA with Tukey’s multiple comparisons test (**B-E**); two-way ANOVA with Tukey’s multiple comparisons test (**F-I**). \**p*<0.05; \*\**p*<0.01; \*\*\**p*<0.001; \*\*\*\**p*<0.0001; ns, not significant.

We next tested whether mTORC1 signaling in sensory neurons is required for this process. Pharmacological inhibition of mTORC1 with sirolimus during the initial priming phase abolished papain-to-HDM cross-sensitization, as indicated by significantly reduced Th2 polarization and cytokine secretion (**Figure 7F-G; Supplementary Figure 6**). Likewise, papain-primed *Scn10a*^Cre/+^*Tfam*^fl/+^ mice failed to mount an enhanced Th2 response to HDM challenge, using both total and HDM-specific readouts (**Figure 7H-I**). Together, these findings demonstrate that mTORC1-driven mitochondrial adaptation in sensory neurons underlies cross-sensitization between antigenically distinct protease allergens, providing a neuronal mechanism for the progressive accumulation of allergen sensitivities characteristic of atopic disease.

## DISCUSSION

Environmental allergens are most often encountered through repeated low-dose exposures, yet how these individually subthreshold encounters cumulatively trigger allergic immunity has remained unclear. Here we identify a sensory neuron-intrinsic mechanism of innate neuroimmune memory, whereby initial activation by protease allergens induces sustained mTORC1 activation, mitochondrial remodeling, and metabolic reprogramming. This primed state renders neurons hyperresponsive to allergens, amplifying itch, enhancing CD301b⁺ DC migration, and augmenting Th2 priming upon re-exposure. Crucially, this mechanism generalizes across antigenically distinct protease allergens, offering a neuronal basis for polysensitization in atopic disease. Our findings also place sensory neurons alongside innate immune cells as potential loci of “trained immunity.”

Analogous to hyperalgesic priming in pain pathways, allergen exposure imprints neurons with a heightened excitability that persists beyond the initial encounter (Reichling and Levine, 2009; Willemen *et al*., 2023). This neuroadaptive response is specifically driven by protease activity. Inert proteins such as Ova fail to induce training, reinforcing the notion that the functional activity of allergens is the trigger for their recognition. The sensors of this functional activity remain undefined. Protease-activated receptors (PARs), such as PAR1, have been linked to neuronal detection of bacterial serine proteases and may play a role in atopic dermatitis pathogenesis (Deng et al., 2023). The HDM cysteine protease Der p 1 cleaves PAR2, generating a non-canonical peptide that may activate PAR2 and the MRGPR family of receptors to promote the release of inflammatory cytokines (Meloun and Leon, 2023; Reddy and Lerner, 2017). However, while PAR2 is cleaved by HDM, it is not required for the activation of sensory neurons (Serhan *et al*., 2019). Alternatively, it is possible that protease-induced stress, through cleavage of membrane proteins, ion influx through membrane damage, or ER stress feeds into mTORC1 activation. Although chronic ER stress canonically inhibits mTORC1 activity, acute activation of ER stress activates mTORC1 activity via ATF6 (Hofmann et al., 2023). This mechanism could explain mTORC1 activation, but how it is maintained in neurons, and whether this maintenance requires metabolic, or growth factor support from surrounding cells such as satellite glial cells remains to be determined (Andreeva et al., 2022). Dissecting these upstream pathways is an important next step.

We show that allergen-induced neuroimmune training is neuron-intrinsic, independent of adaptive lymphocytes, mast cells, or systemic mediators. While resident cells such as keratinocytes or macrophages may contribute, their role is likely secondary. Acute silencing of sensory neurons at the time of allergen exposure abolished training, underscoring neurons as an indispensable gatekeeper of this process, which is mediated through sustained mTORC1 activation. mTORC1 inhibition in Nav1.8^+^ nociceptive neurons prevented training, while persistent activation increased neuronal responsiveness. These data highlight mTORC1 as a molecular switch determining whether allergen encounters are “remembered” by sensory neurons. Mechanistically, mTORC1-mediated memory acts at least in part through PGC-1α-dependent mitochondrial remodeling, which increases OXPHOS capacity without altering glycolysis. Allergen-primed neurons exhibited larger and more interconnected mitochondria, increased oxygen consumption, and enhanced reserve capacity, characteristics associated with robust energy generation. Disruption of mitochondrial stability through *Tfam* haploinsufficiency eliminated training but left acute responses intact, demonstrating that mitochondrial fitness is required for the trained state. These metabolic adaptations likely provide the energetic support required for augmented SP release (a proxy for neuronal activity), which in turn drives CD301b^+^ DC recruitment and Th2 priming. Whether increased mitochondrial output directly leads to greater vesicular release (through ATP availability, calcium handling, etc.) will be an important subject for future investigation, but the link between metabolism and heightened neuronal function is clear.

A striking feature of this pathway is its generality across allergens. Papain priming enhanced neuroimmune responses to HDM, despite lack of antigenic overlap, establishing protease activity as the shared denominator of cross-sensitization. Because both are cysteine proteases, it remains to be tested whether cross-class priming extends to serine or fungal proteases. Nevertheless, this antigen-independent mechanism parallels heterologous memory in innate immune cells and provides a plausible explanation for the clinical progression from monosensitization to polysensitization. Importantly, cross-sensitization required both mTORC1 signaling and mitochondrial stability, identifying them as tractable therapeutic targets. However, the temporal boundaries of the phenomenon remain to be determined. We examined training at a seven-day interval, but its persistence over longer periods remains unknown. Trained immunity in myeloid cells and memory phenotypes in astrocytes involve epigenetic reprogramming (Fanucchi *et al*., 2021; Lee *et al*., 2024; Netea *et al*., 2020; Ochando *et al*., 2023); sensory neurons may similarly acquire chromatin changes that stabilize metabolic and transcriptional states. Given the long lifespan of neurons, even transient changes could have lasting effects, but chromatin changes could act to “lock in” an altered phenotype. Future work will be required to determine whether such epigenetic memory underlies durable neuronal training. In addition, we focused on early immune outcomes – enhanced itch, DC migration, and Th2 priming. Whether neuronal training accelerates later features of allergic disease, including IgE production, mast cell activation, or overt anaphylaxis, remains to be tested.

Together, our results establish mTORC1-driven metabolic adaptation in sensory neurons as a novel form of trained immunity that amplifies allergic sensitization. By shifting the focus from immune-centric to neuro-immune-metabolic mechanisms, this work reframes our knowledge of how allergens initiate and broaden allergic responses. Future studies should identify the upstream sensors of protease activity, define the neuronal subsets most responsible, and explore the durability of this trained state. Targeting sensory neuronal mTORC1 signaling or mitochondrial fitness may provide new strategies to prevent polysensitization and halt the atopic march.

## Limitations of the study

This study demonstrates that protease allergens induce a form of neuroimmune training in sensory neurons through mTORC1-dependent mitochondrial remodeling, but several limitations should be noted. First, the precise upstream mechanisms by which protease activity engages mTORC1 activation in neurons remain undefined. While our data suggest a role for neuronal ion channel activation, the specific molecular sensing of protease allergens requires further study. Second, we primarily assessed training over a 7-day interval; the durability of this state and whether it involves stable epigenetic reprogramming of neurons remain open questions. Third, our model focuses on the initiation of allergic immunity – augmented itch, DC migration, and Th2 priming – without directly addressing later manifestations such as IgE production or mast cell activation. However, these phenomena are presumably downstream of the enhanced Th2 priming that we observed. Finally, cross-sensitization was demonstrated between cysteine proteases, but whether this phenomenon extends across other protease classes or to non-protease allergens remains to be tested. Despite these limitations, our findings reveal a neuron-intrinsic mechanism of neuroimmune memory, providing a framework for understanding the progression from allergen exposure to polysensitization in atopic disease.

## Methods

### Mice

Experiments involving animals were approved by the Massachusetts General Hospital or Harvard Medical School Institutional Animal Care and Use Committee. Mice were maintained and bred in a SPF facility in ventilated cages, with a maximum of five adult mice per cage, on a 12-hour day-night cycle, at 20-25 °C and 35-65% humidity and provided food and water *ad libitum*. All experimental mice were on the C57BL/6 background and were 5-16 weeks of age. Male and female mice were used in all experiments, and comparisons were made between age-and sex-matched controls. Power calculations were conducted to determine sample sizes. Mice were placed into groups depending on their genotype; when possible, mice were randomized. Behavioral experiments were blinded, and immunologic experiments were not. WT C57BL/6 (556) mice were purchased from Charles River Laboratories. WT C57BL/6 (000664), *Rag2^-/-^* (B6.Cg-*Rag2*^tm1.1Cgn^/J; 008449), *Tcra*^-/-^ (B6.129S2-*Tcra*^tm1Mom^/J;002116), *μ*MT^-/-^(B6.129S2-*Ighm*t^m1Cgn^/J; 002288), *Scn10a*^Cre^(B6.129(Cg)-*Scn10a*^tm2(cre)Jwo^/TjpJ; 036564), *Raptor*^fl/fl^(B6.Cg-*Rptor*^tm1.1Dmsa^/J; 013188), *Tsc1*^fl/+^(B6.129S4(Cg)-*Tsc1*^tm1Djk^/MhnyJ; 038428) and *Tfam*^fl/fl^(B6.Cg-*Tfam*^tm1.1Ncdl^/J; 026123) mice were purchased from The Jackson Laboratory. *Cpa3*^Cre/+^ mice were originally from Hans-Reimer Rodewald (German Cancer Research Center). *Rictor*^fl/fl^ mice, originally described by Magee JA, et al. (Magee *et al*., 2012), were generously provided by the laboratory of Dr. Leonid Pobezinsky (University of Massachusetts, Amherst, MA, USA). *Scn10a*^Cre^ were crossed to *Raptor*^fl/fl^, *Rictor*^fl/fl^, *Tsc1*^fl/+^, *Tfam*^fl/+^ to generate *Scn10a*^Cre/+^*Raptor*^fl/fl^, *Scn10a*^Cre/+^*Rictor*^fl/fl^, *Scn10a*^Cre/+^*Tsc1*^fl/+^ and *Scn10a*^Cre/+^*Tfam*^fl/+^ respectively. Kaede mice were originally from Osami Kanagawa as described previously (Perner *et al*., 2020). To control for differences in commensal microbes among colonies, all experiments using *Cpa3*^Cre/+^, *Scn10a*^Cre/+^, *Scn10a*^Cre/+^*Raptor*^fl/fl^, *Scn10a*^Cre/+^*Rictor*^fl/fl^, *Scn10a*^Cre/+^*Tsc1*^fl/+^ and *Scn10a*^Cre/+^*Tfam*^fl/+^ mice were performed with littermate controls bred in our facility. All experiments comparing WT to experimental mouse lines were performed with colony control WT mice originally from The Jackson Laboratory but bred in our facility. In experiments that only used WT mice, the mice were purchased from Charles River Laboratories and housed in the MGH animal facility for one to two weeks prior to undergoing behavioral testing. Mice delivered from animal suppliers not designated for behavioral experiments were used after their arrival to our facility.

### Experimental mouse models

#### Intradermal Immunizations

Mice were briefly anesthetized and injected intradermally (i.d.) with 25 μL of the indicated stimulus. Injections were typically administered into the right cheek except for injections into the left cheek for behavioral testing. The plantar surface of the footpad or the dorsal surface of the footpad (Kaede experiments) was injected for DC migration and/or T cell differentiation. Injections included 1% (w/v) QX314 (Tocris); Papain (50 μg previously frozen, 10 μg freshly prepared (Millipore Sigma), 10 μg freshly prepared (Creative Enzymes), 1μg, 2.5 μg or 5 μg freshly prepared (Fisher Scientific) to normalize for differences in enzymatic activity); heat inactivated papain (50 μg, Millipore Sigma) and Ovalbumin (Ova, Millipore Sigma). For DC migration and T cell differentiation in the multiple low dose exposure model, 1 μg of freshly prepared papain (Fisher Scientific) or 1 μg of house dust mite (HDM) (Greer Laboratories, XPB91D3A2.5) was administered on days 0-2, followed by a 5 μg challenge of HDM on day 9. For behavioral testing in the multiple low dose exposure model, 2 μg of freshly prepared papain (Fisher Scientific) or 5 μg of HDM was administered on days 0 to 2, followed by a 100 μg challenge of HDM on day 9. All reagents were diluted in sterile PBS (Corning). For experiments involving trigeminal ganglia (TG) neuron collection, mice that underwent cheek immunization were anesthetized with an intraperitoneal injection (i.p.) of ketamine (100 mg/kg) and xylazine (10 mg/kg) prior to allergen administration to minimize potential confounding effects of itch and acute inflammation.

#### Behavioral Analysis

As described previously (Flayer *et al*., 2024; Perner *et al*., 2020), mice were transferred from the animal facility to a private behavioral testing room where they were separated into individual cages and underwent a two-hour habituation period. A white noise machine (Marpac) was used in the testing room to reduce distractions. After habituation, mice received an i.d injection of 25 μL of the indicated stimulus into the cheek. Behavioral responses were recorded via overhead video, and the number of scratching bouts directed at the injection site using the hind paw was quantified. A 20-, 30-, or 40-minute video was recorded for papain, and a 25-minute video was recorded for HDM, as indicated in each of the behavior-related figures.

#### Inhibitor Treatment

QX314 (Tocris) was dissolved in PBS (Corning) at 10% (w/v) and stored at −20 °C for up to 3 months. 1% QX314 was co-injected with papain i.d. into the cheek or footpad to assess behavioral responses or DC migration. A-803467 (Selleck) was dissolved in corn oil containing 5% DMSO (Sigma-Aldrich) to a final concentration of 5 mg/mL and stored at 4 °C for up to 10 hours. Mice received i.p injections of A-803467 (50 mg/kg) 4 hours prior to papain exposure for behavioral testing and dendritic cell (DC) migration assays. Sirolimus (StemCell Technologies) and temsirolimus (Sigma-Aldrich) were dissolved in DMSO to prepare 10 mM stock solutions and stored at −20 °C for up to 3 months. For *in vitro* experiments, dissociated dorsal root ganglia (DRG) cultures were treated with sirolimus at a final concentration of 200 nM. For *in vivo* assays, sirolimus (15 nmol in 25 μL PBS) or temsirolimus (12.5 nmol in 25 μL PBS) was administered via i.d. injection in the cheek (behavioral analysis) or footpad (DC migration) 2 hours before papain or sham injections as indicated.

#### DC migration

Mice were briefly anesthetized with isoflurane and injected i.d. in the dorsal footpad with 25 μL of the indicated allergen or 30 μL of TG neuron culture supernatant. For internal controls, one foot received ovalbumin (Ova) while the other received Ova plus indicated allergens in initial and secondary injections. In experiments using Kaede transgenic mice, which express a photoconvertible protein that changes from green (Kaede^Green^) to red (Kaede^Red^) upon exposure to violet light (Tomura *et al*., 2008), DC migration was tracked post-photoconversion. Mice were anesthetized with ketamine (100 mg/kg) and xylazine (10 mg/kg), and the dorsal surface of the feet was exposed to 420 nm light (Bluewave LED curing unit with bandpass filter; Dymax/Andover Corp) for 5 minutes at a distance of 7.5 cm. Allergen was injected i.d. into the photoconverted area immediately afterward. Then, 22-24 hours after allergen injection, popliteal dLNs were collected and digested at 37 °C in RPMI containing DNase I (100 μg/mL; Roche), Dispase II (800 μg/mL; Millipore Sigma), Collagenase P (200 μg/mL; Millipore Sigma), and 1% FBS. Every 7 min, the digestion buffer was removed and transferred into RPMI supplemented with 2 mM EDTA and 1% FBS on ice. Fresh digestion buffer was added to the LNs. Tubes were vigorously shaken before and after each addition of fresh digestion buffer. This process was repeated until no fragment of LN tissue remained. Single-cell suspensions were filtered through a 70 μm cell strainer (Corning or Biologix) before flow cytometry was performed.

#### T cell differentiation

Mice received 25 μL i.d. injections into both dorsal footpads with Ova or Ova co-administered with the indicated allergens. 5 days after the last injections, the popliteal dLNs were harvested and mechanically dissociated in PBS using the frosted end of a Superfrost Plus microscope slide and curved forceps. Single-cell suspensions were filtered through a 70 μm cell strainer (Corning or Biologix). A total of 2 × 10^6^ lymph node cells were resuspended in 1 mL of fully supplemented RPMI medium supplemented with PMA (50 ng/mL; Millipore Sigma), ionomycin (500 ng/mL; Millipore Sigma), and protein transport inhibitor (1:1,000 dilution; BD Biosciences). Cells were incubated for 2.5 to 3 hours at 37 °C with 5% CO₂. Following stimulation, cells were washed twice with FACS buffer and stained per our flow cytometry protocol described below. Additionally, cells were fixed with 4% paraformaldehyde (PFA) for 8 minutes at room temperature (RT) and permeabilized using 1× eBioscience Permeabilization Buffer. Intracellular cytokine staining was performed using fluorescently labeled antibodies for 1 hour at RT or overnight at 4 °C. After two final washes with FACS buffer, samples were analyzed by flow cytometry.

#### Flow cytometry

Single-cell suspensions were prepared by enzymatic digestion and stained in FACS Buffer (1% FBS in DPBS) with fluorescently conjugated antibodies for 30 minutes at 4°C. To ensure viability assessment, Fixable Viability Dye eFluor™ 780 (eBioscience) was added into antibody preparations at a 1:2,000 dilution. Fc receptor blocking was performed using TruStain fcX™ (anti-mouse CD16/32; BioLegend) at a 1:250 dilution. Following staining, cells were washed twice with FACS Buffer to remove unbound antibodies. Flow cytometry analysis was performed using either a CytoFLEX S flow cytometer (Beckman Coulter) or a Cytek Aurora spectral cytometer (Cytek Biosciences). Data acquisition was conducted using manufacturer-specific software (CytExpert v2.3 for CytoFLEX or SpectroFlo v3.2.1 for Aurora). All flow cytometry data (.fcs files) were subsequently analyzed using FlowJo software (v10, TreeStar) with appropriate gating strategies to identify specific cell populations.

#### Isolation of DRG/TG neurons

DRG or TG neurons were isolated from mice as previously described (Flayer *et al*., 2024; Perner and Sokol, 2021). Briefly, tissues were placed in 15 mL Falcon tubes (Corning) containing DMEM (Gibco) supplemented with 10% fetal bovine serum (FBS) and 1% penicillin-streptomycin (Millipore Sigma). Samples were centrifuged at 1,000 rpm for 5 minutes and digested in 3 mL of enzyme solution containing Collagenase A (1.25 mg/mL) and Dispase II (2.5 mg/mL) at 37 °C for 70-90 minutes with shaking at 150 rpm. Following digestion, 10 mL of DMEM with 10% FBS and 1% penicillin-streptomycin was added. Cells were washed twice with DPBS containing 1% penicillin-streptomycin and 1% GlutaMax (Gibco), then resuspended in 1 mL of Neurobasal Medium A (NBM; Thermo Fisher Scientific) supplemented with 1× B-27 and 1% penicillin-streptomycin. Tissues were gently dissociated by pipetting with a P1000 for 10 minutes, then filtered through a 70 μm cell strainer (Corning or Biologix). The resulting suspension was layered onto a 28%/12.5% Percoll gradient (VWR) prepared in Leibovitz’s L-15 Medium (Thermo Scientific) and centrifuged at 1,300 × g for 10 minutes at RT without acceleration or braking. The cell pellet was resuspended in fresh supplemented NBM and filtered through a 70 μm cell strainer (Corning or Biologix) for downstream use.

#### Culture and stimulation of DRG/TG neurons

To prepare the culture substrate, 50 µL of a poly-D-lysine and laminin solution (Millipore Sigma) was added to each well of a tissue-culture-treated, flat-bottom 96-well plate (Corning) and incubated for at least 2 hours at 37°C. The wells were then washed three times with DPBS. DRG or TG neurons (5,000–10,000 cells/well) were cultured for 7 days in supplemented NBM containing 25 ng/mL nerve growth factor (NGF; Alomone Laboratories), 2 ng/mL glial cell line-derived neurotrophic factor (GDNF; PeproTech), and 0.01 mM cytosine β-D-arabinofuranoside hydrochloride (Ara-C; Millipore Sigma). For experiments detecting pS6 and pAKT, the culture medium was replaced with 200 µL/well of supplemented Neurobasal-A medium lacking exogenous growth factors. DRG neuron cultures were either left unstimulated, stimulated with papain (10 mg/mL; Sigma-Aldrich) for 10 min, 30 min, or 1 h, or co-treated with papain (10 mg/mL) and sirolimus (200 nM; StemCell Technologies) for 1 h. After stimulation, supernatants were removed, and cells were harvested for Western blot analysis as described in the ***‘*Protein extraction and immunoblotting’** protocol. To assess Substance P release from TG neuron cultures in response to papain, the culture medium was replaced with 200 µL/well of supplemented Neurobasal-A medium lacking exogenous growth factors. TG neurons were either left unstimulated or stimulated with immobilized papain (Thermo Fisher, 10mg/mL) for 10 minutes. Cell-free supernatants were collected, snap-frozen and stored at −80 °C for subsequent substance P quantification (ELISA) and DC migration assays.

#### Protein extraction and immunoblotting

Cells were lysed in RIPA lysis buffer (G Biosciences) with 1 × protease inhibitor (Thermo Scientific). After a 30-minute incubation on ice, cell lysate was centrifuged at 16000 x *g* for 15 minutes at 4°C. The cell lysates were boiled in SDS loading buffer at 95°C for 5 to 10 minutes and proteins were either run on 10-12% SDS-polyacrylamide gels, followed by transferring to PVDF membranes, or the up-layer supernatants were incubated in SDS loading buffer at 75°C for 10 minutes and proteins run on 4-12% precast gels transferred to nitrocellulose membranes. For blocking non-specific binding and antibody incubation, 5% BSA in 1 × TBST (Cell Signaling Technology) was used. After primary antibody addition (Rabbit anti-pS6(Ser235/236), CST (4858S), 1:1000; Rabbit anti-pAKT (Ser473), CST (9271T), 1:1000; Rabbit anti-PGC1a, Abcam (ab313559), 1:500) samples were incubated at 4°C overnight with shaking. Secondary antibody (Peroxidase (HRP) Anti-Rabbit IgG Goat Secondary Antibody, CST (7074P2), 1:8000) incubation occurred at room temperature for 1 hour. Following secondary incubation, the membranes were washed in 1 × TBST three times. Signal was scanned using BIO-RAD (Hercules, CA, USA) ChemiDocTM MP imaging system or Protein Simple (Biotechne) FluorChem M system.

#### Confocal immunofluorescence of intact ear skin

Confocal imaging of sensory nerve fibers was performed as previously described (Flayer *et al*., 2024). Briefly, ear pinnae were collected from euthanized mice, then the whole ear skin was immediately fixed in 4% PFA for 1 hour at RT. To visualize free nerve endings, fixed ear skin was incubated overnight at 4 °C with anti-Tuj1-biotin antibody (1:100; clone TuJ1, MAB1195; R&D Systems). The following day, samples were incubated with streptavidin-PE (1:100; BioLegend, 405204) to visualize staining. Samples were washed three times with DPBS containing 2% heat-inactivated goat serum and 0.2% Triton X-100, with 10-minute incubations per wash. Intact ear skin was then mounted on SuperFrost Plus microscope slides (Fisher Scientific) using ProLong Diamond Antifade Mountant (Invitrogen). Quantification of Tuj1^+^ sensory nerve fiber density was performed using ImageJ software.

#### Immunofluorescence staining of TOM20 and confocal imaging analyses

Coverslips were circled with an ImmEdge® Hydrophobic Barrier PAP Pen (Vector Laboratories) and coated with 200 µL of poly-D-lysine/laminin solution (Millipore Sigma). Coated coverslips were incubated for ≥2 h at 37 °C and subsequently washed three times with DPBS. TG neurons were isolated from mice 7 days after i.d cheek injections as indicated, following the “**TG/DRG neuron isolation”** method. Approximately 20,000-30,000 neurons were seeded per coated circle and maintained overnight in supplemented NBM containing 25 ng/mL nerve growth factor (NGF; Alomone Laboratories), 2 ng/mL glial cell line-derived neurotrophic factor (GDNF; PeproTech), and 0.01 mM cytosine β-D-arabinofuranoside hydrochloride (Ara-C; Millipore Sigma). For TOM20 staining, cells were fixed with 4% PFA for 8-10 minutes at room temperature and washed three times with DPBS. Cells were incubated with anti-TOM20 antibody (#PA5-52843, Thermo Fisher Scientific; 1:200 in 1% BSA) overnight at 4 °C in a humidified chamber, protected from light. Following three DPBS washes, cells were incubated with the Goat Anti-Rabbit IgG H&L (Alexa Fluor® 647) (ab150079, Abcam; 1:400 in 1% BSA) or at room temperature for 1 hour in the dark. Coverslips were mounted with ProLong Glass Antifade Mountant containing DAPI (Invitrogen, Thermo Fisher Scientific) and stored protected from light. Confocal images were acquired on a Leica Stellaris system equipped with a Power HyD detector, using a 63× oil immersion objective under identical imaging settings for all groups. Mitochondrial morphology quantification was conducted by the Mitochondrial Analyzer Plugin through Image J software (Chaudhry *et al*., 2020). In brief, 3-4 different regions of the image stack were selected, and an average intensity projection of two subsequent stacks per region were calculated. Brightness and contrast were then adjusted to 30-255. Mitochondrial thresholding was then performed in 2D by the Mitochondrial Analyzer, with block size 1.05 and a C-Value of 9 with default settings. Particles lower than 30-pixel units were then removed and “fill holes” was applied. Analysis was then performed with Mitochondrial Analyzer on a per mitochondria basis.

#### Isolation of Genomic DNA from TG neurons

Trigeminal ganglia (TG) neurons were isolated following the “**Isolation of DRG/TG neurons**” protocol. Cells were lysed in 500 µL of lysis buffer (20 mM Tris-HCl, pH 7.4; 100 mM NaCl; 10 mM EDTA; 0.5% SDS) supplemented with 100 µg/mL proteinase K and incubated at 37°C for 30 min. Subsequently, 300 µL of supersaturated NaCl solution was added, mixed thoroughly, and the samples were incubated on ice for 30 min. The lysates were centrifuged at 12,000 × *g* for 15 minutes at 4°C, and the supernatant (aqueous phase) was carefully transferred to a new 1.5 mL microcentrifuge tube. DNA was precipitated by adding 600 µL of isopropanol, mixed gently, and incubation on ice for 30 minutes, followed by centrifugation at 12,000 × *g* for 10 minutes at 4°C. The supernatant was carefully removed to avoid disturbing the pellet, which was then washed with 500 µL of 70% ethanol and centrifuged at 12,000 × *g* for 5 minutes at 4°C. The pellet was air-dried and resuspended in 100 µL of nuclease-free water. DNA concentration and purity were determined using a NanoDrop spectrophotometer.

#### Quantitative PCR (qPCR) and quantification of mitochondrial DNA (mtDNA) copy number

Genomic DNA extracted from TG cells was first diluted with double-distilled water to a final concentration of 10 ng/µL for qPCR analysis. qPCR was performed for the mitochondrial genes 16S ribosomal RNA gene (16S rRNA), NADH dehydrogenase 1 (ND1), and the nuclear-encoded housekeeping gene Hexokinase 2 (HK2) using SYBR Green detection (FastStart Essential DNA Green Master Mix, Roche) on a LightCycler 96 Instrument (Roche). The following primers were used:

HK2: GCCAGCCTCTCCTGATTTTAGTGT (forward) and GGGAACACAAAAGACCTCTTCTGG (reverse);

16S rRNA: CCGCAAGGGAAAGATGAAAGAC (forward) and TCGTTTGGTTTCGGGGTTTC (reverse);

ND1: CTAGCAGAAACAAACCGGGC (forward) and CCGGCTGCGTATTCTACGTT (reverse).

Relative mtDNA copy number was determined by comparing the expression of ND1 and 16S rRNA to that of the nuclear DNA (nDNA) reference gene HK2. The mtDNA copy number was calculated using the following formulas:

ΔCt = Ct (nDNA gene) – Ct (mtDNA gene); Copies of mtDNA = 2 × 2^ΔCt^.

#### Oxygen Consumption Rate (OCR) and Extracellular Acidification Rate (ECAR) measurements of TG neurons

TG neurons were isolated as described in the **‘Isolation of DRG/TG Cells’** protocol. Cells (4000∼5,000 cells/well) were seeded in 8-well Seahorse XFp Cell Culture Miniplates and cultured for 24 hours in supplemented Neurobasal Medium (NBM) under conditions outlined in the **‘Culture and Stimulation of DRG/TG neurons’** protocol. Prior to the assay, cells were gently washed three times and equilibrated in serum-free, unbuffered Seahorse XF RPMI medium (Agilent Technologies) supplemented with 10 mM glucose, 2 mM glutamine, and 1 mM sodium pyruvate. The plate was then incubated at 37°C in a non-CO₂ environment for 60 minutes to allow temperature and pH stabilization. Mitochondrial function was assessed using the Seahorse XFp Cell Mito Stress Test Kit (Agilent Technologies). Sequential injections of metabolic modulators were performed: Oligomycin (5 µM), ATP synthase inhibitor (to assess ATP-linked respiration); FCCP (2 µM), Mitochondrial uncoupler (to induce maximal respiration); Rotenone + Antimycin A (2 µM), then Complex I/III inhibitors (to determine non-mitochondrial respiration). Metabolic Parameter Calculations were derived as follows:

1. Basal respiration = [OCR (basal)] – [OCR (Rotenone/Antimycin A)].
2. ATP-linked respiration = [OCR (basal)] – [OCR(Oligomycin)].
3. Maximal respiratory capacity = [OCR (FCCP-induced peak)] – [OCR (Rotenone/Antimycin A)].
4. Spare respiratory capacity = [OCR (FCCP-induced peak)] – [OCR (basal)].
5. Coupling efficiency = ATP-linked respiration / Basal respiration.

#### *In vitro* stimulation of draining lymphocytes

Draining lymphocytes enzymatically digested from the dLN after completing the ‘**DC migration**’ protocol were added to a 96 well flat bottom plate (Corning) at 5×10^5^ cells/per well. 30µg or 100µg of heat inactivated HDM (HI-HDM) in 200 µL of fully supplemented RPMI was added to the cells. Plates were incubated at 37 °C with 5% CO_2_ for 4-5 days until the medium turned yellow, then cell-free supernatant was harvested, snap-frozen and stored at −80 °C for downstream analysis.

#### Enzyme-Linked Immunosorbent Assay (ELISA)

Substance P levels were quantified using a competitive mouse Substance P ELISA kit (Abcam) following the manufacturer’s protocol. For cytokine analysis, cell-free culture supernatant from stimulated whole dLN cells was collected. High-binding 96 well plates (Corning) were coated with capture antibodies overnight at 4 °C. IL-4, and IL-13 were captured with anti-mouse IL-4 (0.25 mg/ml, BioLegend) and anti-mouse IL-13 (1.0 mg/ml, BioLegend), respectively. Plates were then washed three times with 0.1% Tween 20 in PBS and blocked with 1% (wt/vol) BSA for 2 hours at 37 °C before cell-free culture supernatant was plated overnight at 4 °C. Next, plates were washed three times with 0.1% Tween 20 in PBS. IL-4 and IL-13 were detected by adding biotinylated anti-mouse IL-4 (0.25 mg/ml, BioLegend) or biotinylated anti-mouse IL-13 (0.25 mg/ ml, eBioscience) for 2 hours at RT. Following this, plates were washed four times, incubated with Streptavidin-horseradish peroxidase (BioLegend) for 30 minutes at RT, washed 7 times, and finally incubated with OPD solution (Sigma-Aldrich) for development. Development was stopped with 2M H_2_SO_4_. ELISA samples were assessed on a SpectraMax iD5 microplate reader (Molecular Devices). Concentrations were determined from a standard curve.

#### Bulk RNA-seq data generation

To prepare sequencing libraries, total RNA was extracted from ground TG tissue using TRIzol® reagent (Invitrogen). Then, 20 ng of RNA lysate was processed using a modified SMART-Seq2 protocol (Picelli et al., 2014) entailing RNA secondary structure denaturation (72°C for 3 min), reverse transcription with Maxima Reverse transcription (Life Technologies), and whole-transcriptome amplification (WTA) with KAPA HiFi HotStart ReadyMix 2X (Kapa Biosystems) for 12 cycles. WTA products were purified with Ampure XP beads (Beckman Coulter), quantified with a Qubit double-stranded DNA HS Assay Kit (Thermo Fisher Scientific), and quality assessed with a high-sensitivity DNA chip (Agilent). The purified WTA product (0.2ng) was used as input for the Nextera XT DNA Library Preparation Kit (Illumina). Uniquely barcoded libraries were pooled and sequenced with a NextSeq 500/550 high-output V2 75 cycle kit (Illumina) using 2x 38 paired end reads.

#### Bulk RNA-seq read alignment and quantification

Raw sequencing data was processed using the bulk_rna_seq (xavier-genomics/bulk_rna_seq, v24) workflow from the Broad Institute’s online platform Terra. Reads from each sample were aligned to the mouse reference genome GRCm38 through STAR alignment by using “GRCm38_ens93filt” as the reference and “star” as the aligner. Output expected count matrices were combined to produce one sample x gene matrix.

#### Differential expression analysis

The package DESeq2 (v1.38.2, R) was used to identify changes in gene expression between treatments using gene ∼ treatment as the model design. The raw count matrix was provided as input.

#### Gene set enrichment analysis

GSEA was performed using the fgsea function from the fgsea package (v1.18.0, R) with 10,000 permutations to test for independence. Input gene sets tested were derived from the HALLMARK pathways database and limited to those with a minimum of 10 genes per set. For all comparisons of treatment, input gene rankings were based on decreasing Wald statistic values from DE analysis. The package biomaRt (v2.65.0, R) was used to map mouse genes to their human orthologs by referencing the Ensembl database.

#### Quantification and Statistical analysis

Analysis of data was performed using GraphPad Prism version 10.3.1 (GraphPad Software). Comparison of data between two groups was performed using Welch’s t-tests. Comparison of data between more than two groups with one independent variable was performed using one-way ANOVA with Tukey’s multiple comparisons tests. Comparison of two independent variables was performed using two-way ANOVA with Tukey’s multiple comparisons tests. P-values of less than 0.05 were considered statistically significant. Statistical details are listed in the respective figure legends.

#### Inclusion and ethics statement

Roles and responsibilities were agreed among all authors during the inception of this study and before any work was conducted. Experiments involving animals were approved by the Massachusetts General Hospital or Harvard Medical School Institutional Animal Care and Use Committee.

## Acknowledgments

This work was supported by the National Institute of Allergy and Infectious Diseases (grant R01AI151163 to C.L.S.), the Food Allergy Science Initiative (C.L.S.), a Massachusetts General Hospital Howard M. Goodman Fellowship (C.L.S.), the Gene Lay Institute of Immunology and Inflammation (C.L.S.), the Gene Lay Institute of Immunology and Inflammation Fellowship (X.Z.), and a shared instrumentation grant from the National Institutes of Health (1S10OD036287). We thank Rod A. Rahimi for his valuable advice and mentorship. We are grateful to Melina Sokolowska and Nino Stocker (Swiss Institute of Allergy and Asthma Research, University of Zürich, Davos, Switzerland; Christine Kühne – Center for Allergy Research and Education, Davos, Switzerland) for their insight and guidance on mitochondrial morphology analysis. We thank Leonid Pobezinsky (University of Massachusetts, Amherst) for providing *Rictor*^fl/fl^ mice and Joshua A. Boyce and Hans Reimer Rodewald for providing *Cpa3*^Cre/+^ mice. We also thank members of the Sokol laboratory for their feedback and suggestions. Graphical abstracts were created using BioRender.com. Cytometric analyses were performed with support from the MGH Department of Pathology Flow Cytometry Core.

## Author contributions

Experiments were conceived, designed, performed, and analyzed by X.P.Z. (all), H.B.Y. (Figs. 4B-C, 6A-B; and Supplementary Figs. 4D, 5E, 6A-B), H.T.Z. (Figs. 5H, 7B-D; and Supplementary Figs. 4C, 5F), I.J.K. (Figs. 4A-B; and Supplementary Figs. 4A-B and Table 1-1), N.L. (Figs. 2G-H, 5F-G, 6H), D.R.B. (Figs. 2A-B, 3E, 7B), C.H. (Figs. 3G, 4C), P.R.N. (Figs. 3D, 5A-B) and C.L.S. (all). E.W. assisted with bulk-seq library generation. L.M.A. and P.R.M. assisted with Leica Stellaris system and confocal imaging. N.P.S. assisted with bulk-seq analysis. R.L. assisted with behavioral assays. C.H.F., S.Z., and R.A.R. provided technical and experimental assistance. Z.W.S. assisted with Seahorse analysis. A.-C.V. designed and oversaw bulk-seq sequencing. X.P.Z. and C.L.S. wrote the manuscript. P.R.M., S.Z., C.H., D.R.B., and E.W. contributed to manuscript editing. C.L.S. provided resources, reagents, funding, and supervised the study.

---

**Supplementary Figure 1:**
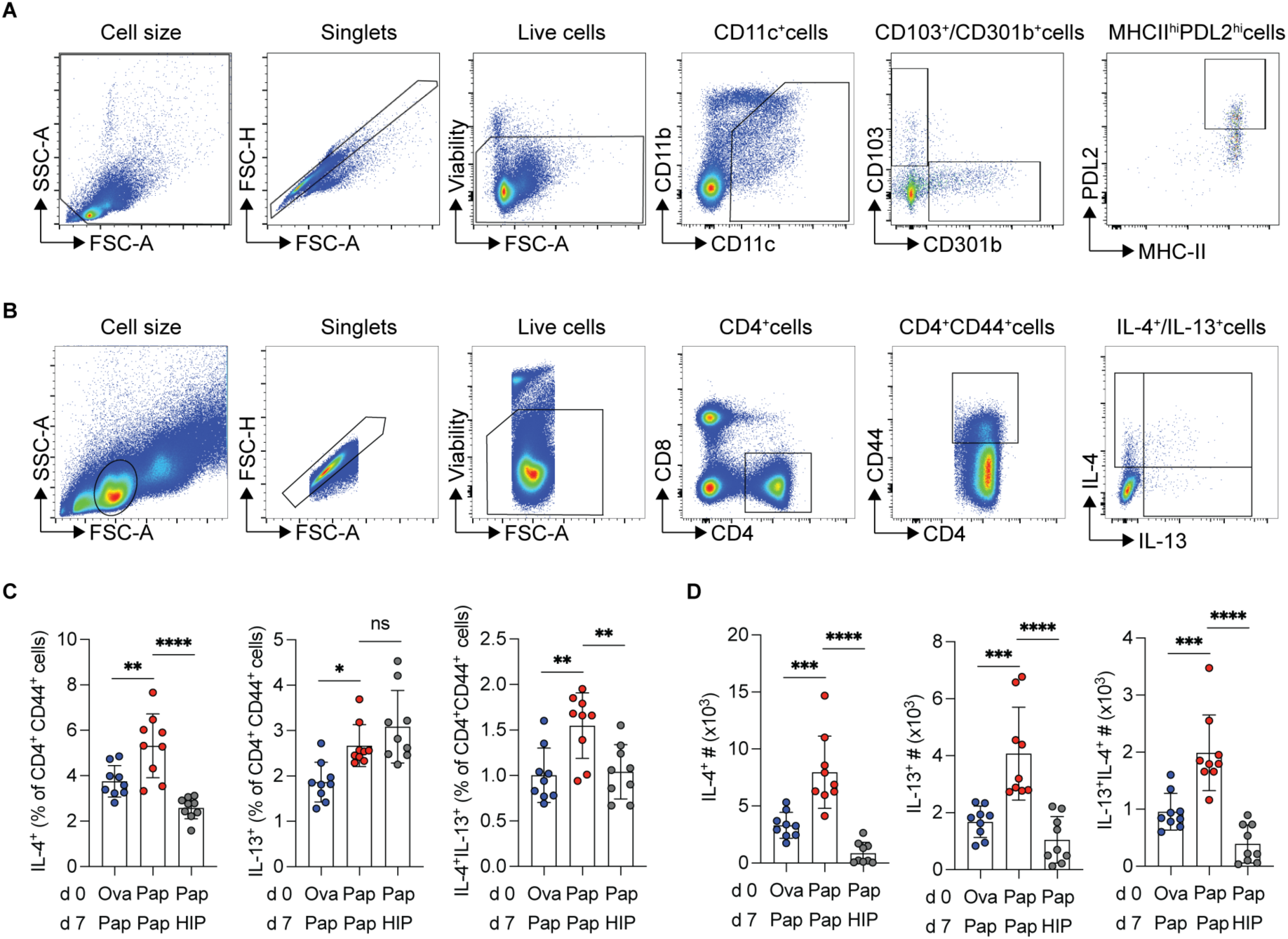
The CD4 T-cell response in protease allergen-induced neuroimmune memory requires protease activity. (**A**) Representative gating strategy for flow cytometric analysis of DCs in popliteal draining lymph node (dLN) from wild type (WT) mice that received intradermal (i.d.) papain footpad injection. (**B**) Representative gating strategy for Th2 cytokine analyses in popliteal dLN from papain footpad-injected WT mice. (**C-D**) Flow cytometric analysis of IL-4⁺, IL-13⁺, and IL-4⁺IL-13⁺ cells within the CD4⁺CD44^high^ T cell population in the popliteal dLN 5 days after secondary injections as indicated, shown as percentage (**C**) and cell number (**D**). Each data point represents an individual mouse. Bar plots are mean ± S.D. All data represent at least two independent experiments with each experiment including ≥3 mice per group. All plots show data combined from multiple experiments. Statistical tests: ordinary one-way ANOVA with Tukey’s multiple comparisons test (**C-D**). \**p*<0.05; \*\**p*<0.01; \*\*\**p*<0.001; \*\*\*\**p*<0.0001; ns, not significant.

**Supplementary Figure 2:**
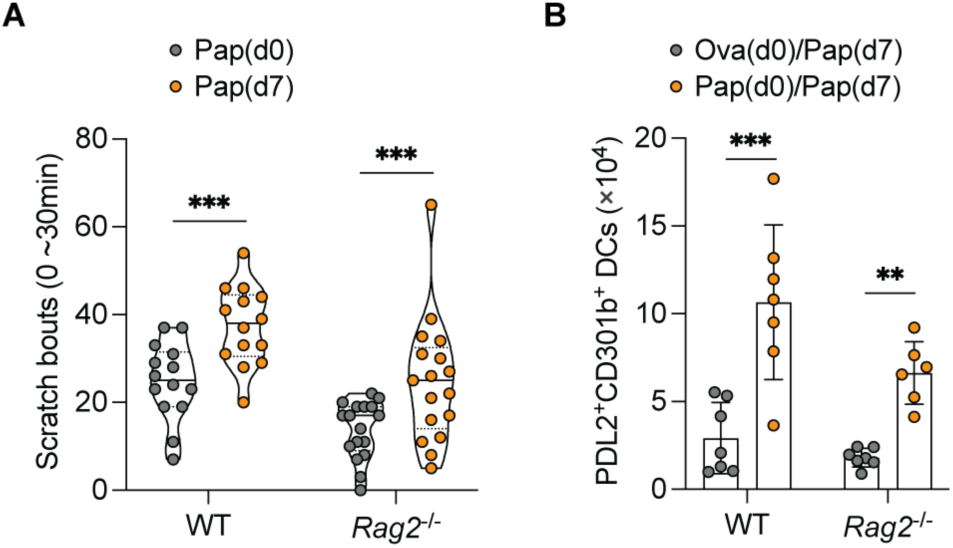
Allergen-induced neuroimmune memory occurs independently of lymphocytes. (**A**) WT or *Rag2^-/-^* mice were i.d. injected with papain into the cheek on days 0 and 7. Scratching behavior was quantified as the total number of scratch bouts at the site of secondary injection. (**B**) Flow cytometric analysis of PDL2⁺CD301b⁺ cells in the popliteal dLN from WT and *Rag2^-/-^* mice 22-24 h after secondary footpad injections as indicated. Each data point represents an individual mouse. Violin plots show the median and quartiles. Bar plots are mean ± S.D. All data represent at least two independent experiments with each experiment including ≥3 mice per group. All plots show data combined from multiple experiments. Statistical tests: two-way ANOVA with Tukey’s multiple comparisons test (**A-B**). \*\**p*<0.01; \*\*\**p*<0.001.

**Supplementary Figure 3:**
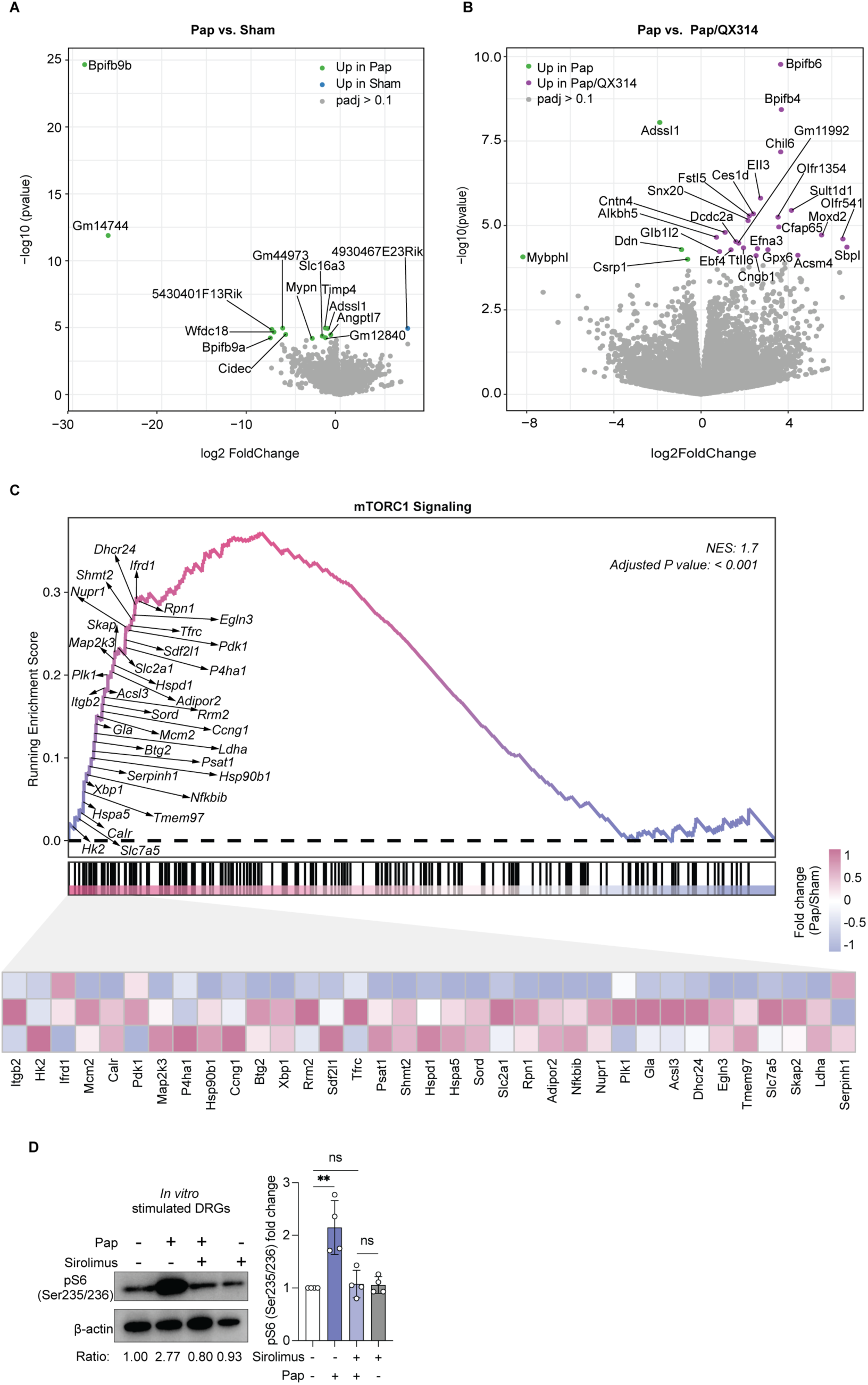
Protease allergen-stimulated sensory neurons exhibit mTORC1 activation. (**A-B**) Volcano plots showing differentially expressed genes in trigeminal ganglia (TG) neurons 7 days after a single i.d. cheek injection with Ova (Sham), papain (Pap), or papain co-administrated with QX314 (Pap/QX314). Pap vs. Sham is shown in (**A**) and Pap vs. Pap/QX314 is shown in (**B**). Volcano plots show –log10(*p* value) versus log2 fold change. Genes considered significant were determined based on adjusted *p* values (padj > 0.1). Differentially expressed genes are labeled in the volcano plot. **(C)** Gene set enrichment analysis (GSEA) reveals the over-representation of the mTORC1 signaling pathway in TG neurons isolated from WT mice as in (**A-B**). The upper part of the plot shows the distribution of the top 35 signature genes involved in the mTORC1 signaling pathway in TG neurons from papain-injected cheeks. The lower heatmap presents the relative expression of these leading-edge genes (Fold change was calculated as the per-gene counts in each papain-exposed TG neuron sample divided by the mean gene counts of OVA-exposed TG neurons). Fold change (FC) values in the heatmap were normalized by row. FDR P-value, two-sided. NES, normalized enrichment score. GSEA was performed based on genes differentially expressed in the Pap group relative to the Sham group. (**D**) Representative immunoblots (left) and quantification (right) of pS6 (Ser235/236) levels in DRG neurons stimulated *in vitro* with papain or papain and sirolimus. The normalized pS6/actin ratio is shown below each lane. Bulk-seq in (**A-C**) was performed with three replicates per group. Bar plots are mean ± S.D. Each dot in the bar plot of (**D**) represents one independent assay. Statistical analysis: two-tailed ratio paired *t*-test (**B**, **C**), \*\**p*<0.01; ns, not significant.

**Supplementary Figure 4:**
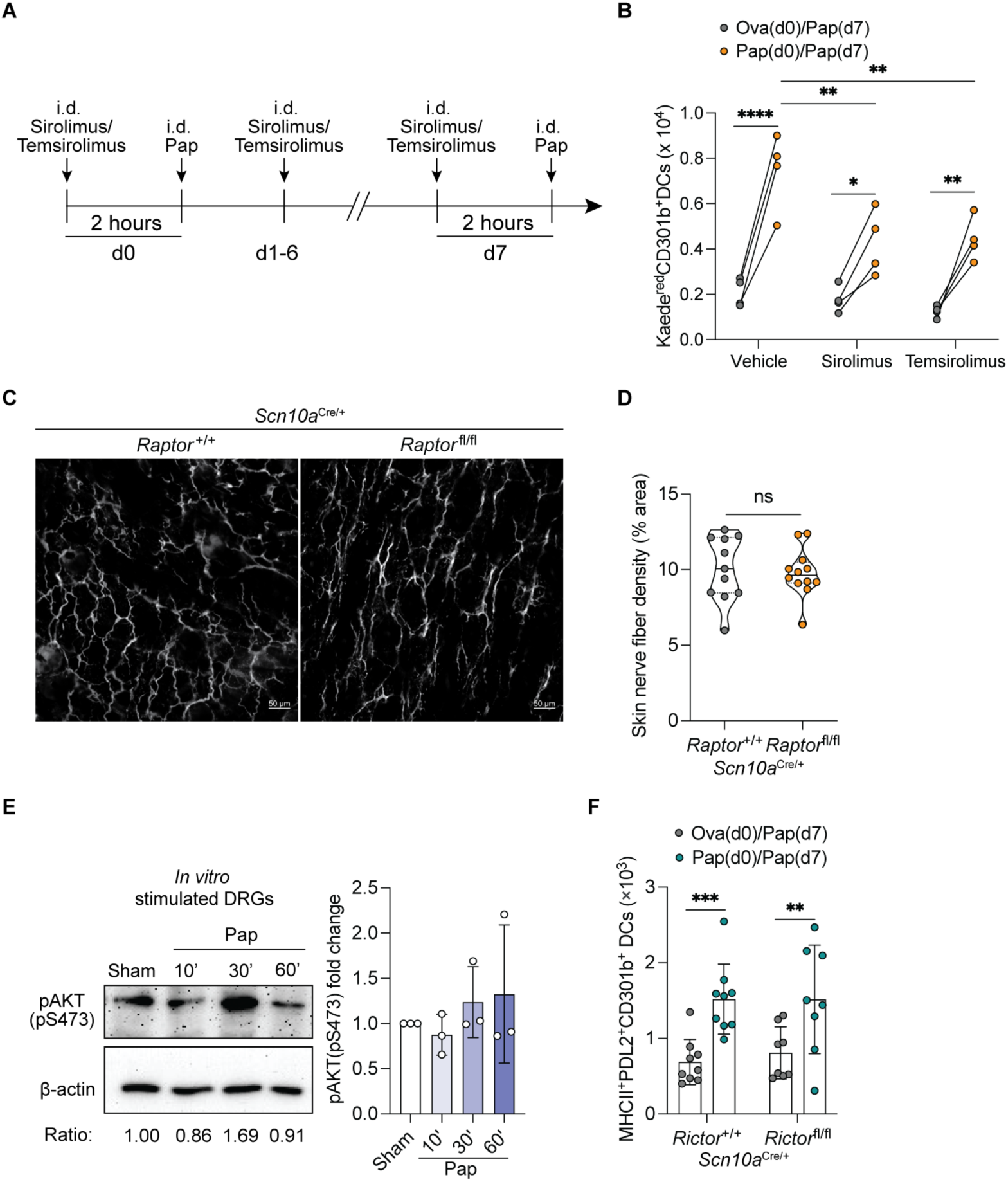
Sensory neuronal mTORC1 activation is required for allergen-induced neuroimmune memory. (**A**) Schematic of sirolimus or temsirolimus treatment protocol for assessing behavior responses and DC migration. Briefly, WT mice received i.d. cheek (for behavioral response) or footpad (for DC migration) injections of sirolimus, temsirolimus, or vehicle 2 hours before papain injection on day 0, followed by daily dosing on days 1-6 and a final dose 2 hours before secondary papain injection on day 7. Scratching behavior was quantified as the total number of scratch bouts at the site of secondary injection. DC migration into the popliteal dLN was tracked 22-24 hours after secondary papain injections as indicated. (**B**) Kaede^GFP^ mice were administered either sirolimus or temsirolimus as shown in (**A**) followed by i.d. footpad injections as indicated on day 0 and then photoconversion at the initial injection site followed by papain injection on day 7. The total number of Kaede^Red^CD301b⁺ DCs 22-24 hours after secondary injections in the popliteal dLN was quantified by flow cytometry. (**C**) Confocal microscopy z-stacks showing Tuj1 expression in the skin of *Scn10a*^Cre/+^*Raptor*^fl/fl^ and littermate control *Scn10a*^Cre/+^*Raptor*^+/+^ mice. The scale bar is 50 μm. (**D**) Violin plots show quantification of skin nerve fiber density. Each data point represents an individual image. (**E**) Representative immunoblots (left) and quantification (right) of pAKT (Ser473) levels in DRG neurons stimulated *in vitro* with papain. The normalized pAKT/actin ratio is shown below each lane. (**F**) Flow cytometric analysis of PDL2⁺CD301b⁺ DCs in the popliteal dLN from *Scn10a*^Cre/+^*Rictor*^fl/fl^ and littermate control *Scn10a*^Cre/+^*Rictor*^+/+^ mice following i.d. footpad injections as indicated. Each data point in (**B, F**) represents an individual mouse. Each dot in the bar plot of (**E**) represents one independent assay. Bar plots are mean ± S.D. Violin plots show the median and quartiles. All data represent at least two independent experiments with each experiment including ≥3 mice per group. All plots show data combined from multiple experiments, except in (**B**) where one representative experiment is shown. Statistical tests: two-way ANOVA with Tukey’s multiple comparisons test (**B, F**); two-tailed unpaired *t*-test (**D**). *p<0.05; **p<0.01; ***p<0.001; ****p<0.0001; ns, not significant.

**Supplementary Figure 5:**
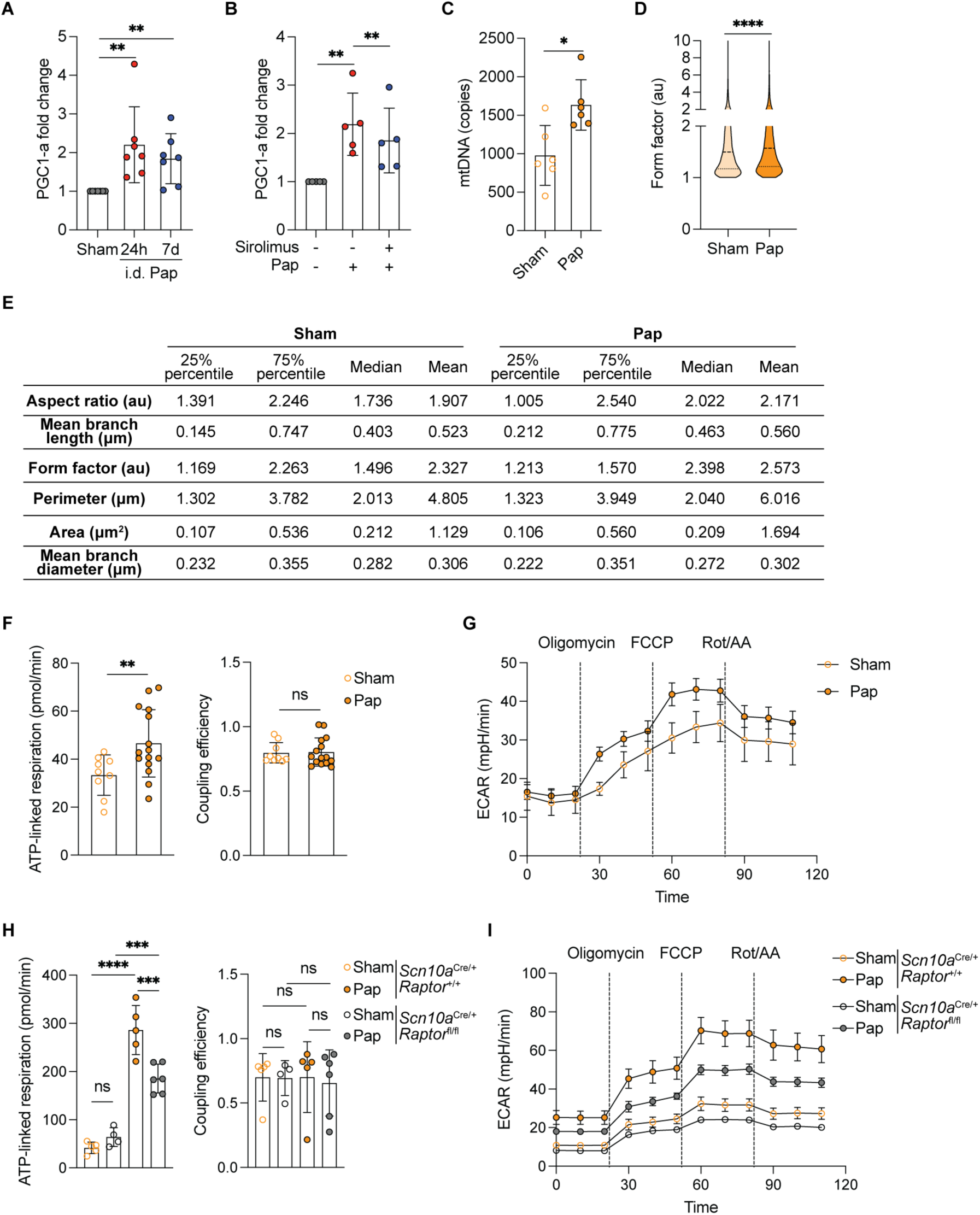
mTORC1 promotes mitochondrial function required for allergen-induced neuroimmune memory. (**A**) Quantification of PGC-1α protein levels in TG neurons from WT mice 24 hours or 7 days after i.d. cheek injection of Ova (Sham) or papain (Pap). (**B**) Quantification of PGC-1α protein levels in TG neurons harvested 7 days after papain injection with or without sirolimus treatment. WT mice received i.d. cheek treatment with vehicle or sirolimus 2 hours before i.d. papain injection on day 0, followed by an additional i.d. cheek treatment on day 1 to ensure complete inhibition of early allergen-induced mTORC1 activation. (**C**) Quantification of mitochondrial DNA (mtDNA) copy number in TG neurons isolated from WT mice 7 days after i.d. cheek injections as indicated. (**D**) Violin plots show quantification of mitochondrial form factor (arbitrary units, au) in sham (∼170 neurons) or papain-primed (∼194 neurons) TG neurons obtained with the Mitochondrial Analyzer plugin in Image J. (**E**) Table shows morphometric analysis of mitochondria in TG neurons isolated as in (**C**). Values represent quartiles (25% percentile and 75% percentile), median, and mean for each parameter quantified using the Mitochondrial Analyzer plugin in image J. (**F**) Quantification of the ATP-linked respiration and coupling efficiency of TG neurons isolated as in (**C**). (**G**) Extracellular Acidification Rate (ECAR) as determined by Seahorse metabolic profiling in response to oligomycin, FCCP (carbonyl cyanide 4-(trifluoromethoxy) phenylhydrazone), Rot/AA (rotenone and antimycin A) of TG neurons isolated as in (**C**). (**H**) Quantification of the ATP-linked respiration and coupling efficiency of TG neurons isolated from *Scn10a*^Cre/+^*Raptor*^fl/fl^ and *Scn10a*^Cre/+^*Raptor*^+/+^ littermate control mice 7 days after i.d. cheek injections as indicated. Sham groups for each genotype are indicated by open circles, and papain groups by filled circles. (**I**) ECAR profiles were determined by Seahorse metabolic profiling as in (**G**) of TG neurons isolated from mice as described in (**H**). Sham groups for each genotype are indicated by open circles, and papain groups by filled circles. Each dot in the bar plot of (**A-B**) represents one independent assay. Each data point in (**C, F-I**) represents an individual mouse. Violin plots show the median and quartiles. Bar plots are mean ± S.D. All data represent at least two independent experiments with each experiment including ≥3 mice per group. All plots show data combined from multiple experiments. Statistical analysis: two-tailed ratio paired *t*-test (**A-B**); two-tailed ratio paired *t*-test un (**C-E**); ordinary one-way ANOVA with Tukey’s multiple comparisons test (**G**). **p<0.01; ***p<0.001; ****p<0.0001; ns, not significant.

**Supplementary Figure 6:**
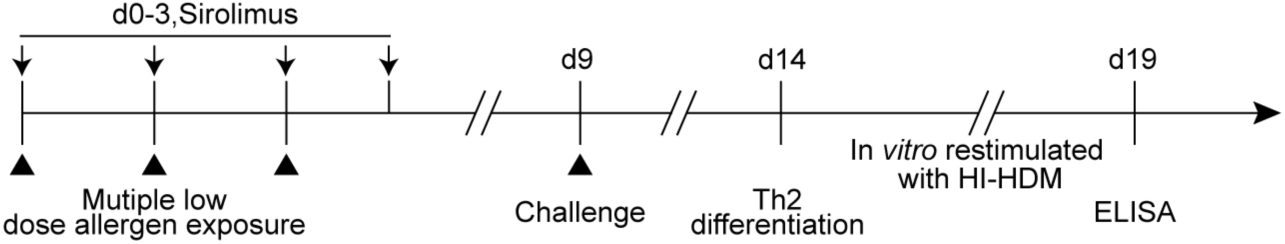
Allergen-induced neuroimmune memory promotes responsiveness to recurrent low dose allergen exposure in an antigen-independent manner. Schematic of sirolimus treatment in multiple low dose allergen exposure model to assess the role of mTORC1 signalling in T cell differentiation and HDM-specific T-cell responses by ELISA. WT mice received i.d. injections of vehicle or sirolimus 2 hours prior to papain (Pap) footpad injections on days 0-2, followed by an additional sirolimus injection on day 3. On day 9, mice were challenged with the indicated allergens. Th2 differentiation in the popliteal dLN was assessed by flow cytometry. Whole dLN cells were cultured *in vitro* with heat-inactivated house dust mite (HI-HDM) for 4-5 days until culture media turned yellow. HDM-specific IL-4 and IL-13 production in culture supernatants was measured by ELISA.

